# Improved Estimation of Bird Abundance and Energy Consumption by Combining Point and Transect Observations

**DOI:** 10.1101/2023.08.21.554181

**Authors:** Maria Jose Ayala Bolagay, Aaron Brothers, Marta Quitián, Eike Lena Neuschulz, Matthias Schleuning, Katrin Böhning-Gaese, Orlando Alvarez, Vinicio Santillán, Christopher Thron

**Affiliations:** Universidad Católica de Cuenca, Ecuador; Texas A&M University-Central Texas, Killeen TX USA; Instituto Mediterráneo de Estudios Avanzados (CSIC-UIB); Senckenberg Biodiversity and Climate Research Centre, Frankfurt Germany

**Keywords:** Energy consumption, Abundance, Estimation, Survey, Point counts, Transect sampling, Bird, Ecosystem, Frugivore, Species, Family, Simulation

## Abstract

Point counts and transect surveys are both common methods for estimating bird abundance (by species, genus, or family) in a given environment. Sometimes both methods are used in the same study. Point counts reflect abundance directly, while transect observations of bird feeding instances reflect both the bird abundance and the mean energy consumption rate per bird.

In this research we develop an improved estimator of bird abundance based on both sets of measurements, and derive error bars for these abundance estimates. The improved estimator is a linear combination of the two individual abundance estimates obtained from the two data types. The combination weights are obtained through a comparison of two separate energy estimates. The mathematical methods developed are applied to a dataset consisting of point counts and transect surveys of Ecuadorian frugivore bird species in six different types of environment (three elevations and two degrees of forestation). The improvement in accuracy achieved by this combined estimator is verified by simulations applied to bird family data. We also obtain 90% confidence intervals for the bird family relative abundance estimates, and verify the confidence intervals using simulations.

## 1 Introduction

Two common methods for estimating abundance of bird types (whether species, genera, or families) in a given area are point counts and transect surveys. Point counts are obtained when the observer chooses a set of fixed observation points within the area of interest, and counts bird detections (by sight or by sound) during multiple sessions over an extended period of several days or months. Transect surveys are conducted by the observer specifying a transect (i.e. narrow section) within the area and repeatedly walking through and recording bird observations at the same time of day over an extended period. Transect measurements can target bird feedings, so that the frequency of feeding on each different type of plant is recorded.

Each of the two survey types has distinct advantages and limitations. Point counts are straightforward to conduct and are effective for detecting a broad cross-section of species at fixed locations, but they do not directly record feeding behavior. Transect surveys, by contrast, capture behavioral data including feeding events and plant-bird interactions, but coverage of the survey area depends on the observer’s path and pace, which can introduce systematic biases. Both methods have been widely adopted because neither alone provides a complete picture of bird community structure and energy dynamics; combining them allows investigators to cross-validate abundance estimates and extract behavioral inferences that neither method supports independently.

The energy consumption of different bird types can be expressed in two different ways. The total energy consumption of a bird type can be expressed as the number of individuals of that type times the mean consumption per individual. On the other hand, the same quantity may be expressed as the total number of feedings by birds of that type times the amount consumed per feeding. Point counts can be used to estimate bird abundance required for the first expression, while transect surveys can provide feeding data for the second expression.

We may make use of these two alternative estimates of energy consumption in several different ways. First, by comparing the two estimates, we may gain information about bird consumption rates. Second, we may combine the two estimates to obtain a more accurate estimate of energy consumption per bird type, and hence of bird type relative abundance. Finally, given that count measurements are Poisson distributed (which is commonly recognized as valid for count measurements), we can use the results obtained and simulations to obtain confidence intervals for the relative abundance of different bird types.

The methodologies that we have developed may be applied to other surveys where multiple forms of data related to species abundance are present. Our results can be used both to obtain more accurate abundance estimates, and to rigorously specify accuracies of the estimates thus obtained. We have also demonstrated a technique for estimating an important biological parameter, namely the exponent in the functional dependence of bird energy consumption on bird weight. Using this technique, other datasets may provide similar estimates.

The remainder of this paper is organized as follows. Section 2 surveys previous similar studies. Section 3 consists of two parts. In the first part, we present the theoretical derivations for the statistical methods used for the data analysis. These methods include the construction of point count-based and transect observation-based estimators for energy consumption and for bird abundance; and the combination of bird abundance estimators and log abundance estimators into more accurate estimators. In the second part, the methods used to evaluate the theoretical results are described. These methods include the use of an empirical dataset obtained from several regions in southern Ecuador; this dataset is used both to estimate model parameters from bird species data, and as a basis for simulations to test the accuracy of the different theoretical estimators using bird family data. In Section 4 the results of the evaluation are presented; all figures displaying these results are placed at the end of the paper, after the bibliography. Section 5 discusses the implications of our results in relation to prior work. Finally, in Section 6 our conclusions from this study are summarized.

## 2 Previous studies

Frugivore bird species play a critical role in tropical and subtropical ecosystems, functioning as primary agents of seed dispersal and thereby enabling plant population regeneration, maintaining plant diversity, and sustaining ecological balance [1]. The ecological importance of these interactions has motivated a substantial body of empirical research across diverse geographic regions. Studies in the Neotropics, Africa, and Asia have documented that frugivore bird assemblages differ markedly in composition and foraging behavior across habitats and elevational gradients, and that the intensity of fruit consumption and seed dispersal is closely tied to local bird abundance [2, 3, 4, 5, 6, 7, 8]. Several of these studies have specifically examined the relationship between bird body size and dietary preferences, noting that larger frugivores tend to consume larger fruit and disperse seeds over greater distances, while smaller species are restricted to smaller fruit and contribute to fine-scale seed rain patterns [7, 8]. Work in Andean systems in particular has highlighted how frugivory and seed dispersal effectiveness vary with elevation and habitat fragmentation, with fragmented landscapes typically supporting depauperate communities of large-bodied frugivores and altered plant-bird interaction networks [6, 5]. Understanding how bird abundance responds to these environmental gradients is therefore essential for predicting the cascading effects of habitat change on ecosystem function.

Accurately quantifying bird abundance in natural environments requires careful choice of census methodology, and the merits of different field approaches have been extensively debated in the literature [9, 10, 11]. Point count methods, in which observers record all detections within a fixed radius or time window from a stationary position, are widely used because they are standardizable, repeatable, and efficient over large spatial extents. However, point counts depend heavily on observer skill and may miss secretive or cryptic species. Transect surveys, in which observers walk a predetermined route and record detections along the way, are better suited for capturing behavioral information such as feeding events and plant-bird interactions, but require careful control of walking pace and can introduce distance-related detection biases. Because the two methods sample different aspects of bird community activity, combining them in a single study can provide richer information than either method alone. This complementarity motivates the analytical framework developed in the present paper, which explicitly links the two data types through a shared model of energy consumption, allowing each to inform and constrain the other.

Statistical methods play an increasingly important role in extracting reliable estimates from field survey data, particularly when sample sizes are modest or data are missing. Bayesian approaches offer a principled framework for incorporating prior biological knowledge—such as the expected dependence of metabolic rate on body mass—into parameter estimation, and for propagating uncertainty through hierarchical models of abundance and detection [12, 13]. The methods proposed in the present paper operate within this general philosophy: by exploiting the theoretical relationship between point counts and transect feeding observations, we derive estimators that borrow strength from both data types, and we use simulation-based bootstrapping to characterize estimation uncertainty without requiring distributional assumptions that may not hold in practice. This approach yields abundance estimates with formally justified confidence intervals, contributing to a more rigorous quantification of bird community structure than is possible from either data source alone.

## 3 Methodology

This section consists of two subsections dealing with theoretical and empirical methods, respectively. The first section derives statistical estimators that make use of point counts and transect observations to estimate model parameters and demographics. The second section describes the application of these theoretical results to a specific ecological survey.

### 3.1 Theoretical equations

#### 3.1.1 Transect observation-based expression for energy consumption per bird type

The energy consumption per bird type over a given period of time at a given location may be expressed as the total number of feedings times the mean energy transfer per feeding. We may rewrite this statement in the form of an equation:

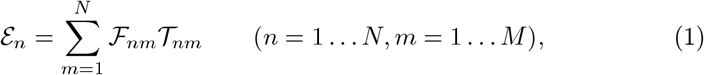

where

- *M, N* : Number of fruit and bird types, respectively.
- ℰ_*n*_: Total energy consumption of all birds of type *n* at the specified location during the specified period.
- ℱ_*nm*_: Total number of feedings of bird type *n* on fruit type *m*; 𝒯_*nm*_: Mean energy transfer per feeding for bird type *n* on fruit type *m*.

Under the assumption that the transect observations are representative and capture the same proportion of all bird-fruit interactions, we have:

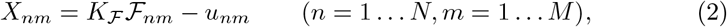

where

- *K*_ℱ_ : Constant of proportionality, assumed to be independent of bird and fruit types;
- *X*_*nm*_: Number of transect observations of bird type *n* consuming fruit type *m*;
- *u*_*nm*_: Observational uncertainty for bird type *n* consuming fruit type *m*.

We furthermore assume initially that energy transfer per feeding depends on both the mean fruit weight and the mean bird weight. This assumption presumes that larger fruit may require less foraging, and larger birds may be able to consume more per feeding. This relationship is summarized in the equation:

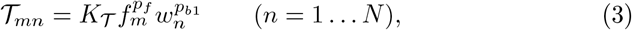

where

- *K*_𝒯_ : Constant of proportionality, assumed to be independent of bird type.
- *f*_*m*_: mean fruit weight for type *m*.
- *w*_*n*_: Mean weight of bird type *n*;
- *p*_*f*_, *p*_*b*1_: Exponents of fruit weight and bird weight respectively, in power model.

Substituting ℱ_*nm*_ and 𝒯_*nm*_ in (1) with their expressions from (2) and (3) respectively, we have finally:

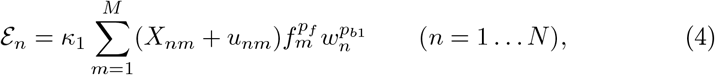

where *κ*_1_ ≡ (*K*_ℱ_ *K*_𝒯_)^−1^ is a constant of proportionality that is independent of bird and fruit types. In order to simplify the model and make it amenable to analysis, we replace *f*_*m*_ in (4) with 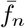, which is defined as the mean fruit weight consumed by bird species *n*. This simplification will be accurate when each bird species mostly consumes fruit species with similar average weights. An additional simplification concerns the magnitude of the measurement errors *u*_*nm*_. We assume that these errors are nearly constant in percentage across species: that is to say, the error for species *n* will be roughly proportional to the number of observations *X*_*n*_.

With the two simplifications described above, the final simplified model is given by the equation:

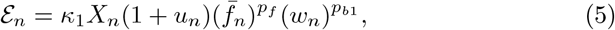

where

- *X*_*n*_: Sum of transect observations for bird type *n*. The formula for *X*_*n*_ is 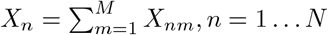.
- *u*_*n*_: Combined proportionate observational uncertainty for bird type *n*. The variables {*u*_*n*_}, *n* = 1, … *N* are modeled as independent, identically distributed random variables with mean 0 and variance 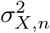;
- 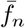: Average weight of fruit consumed by bird type *n*, given by the formula:

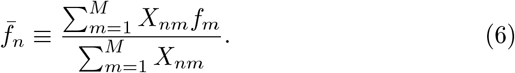

#### 3.1.2 Point count-based estimate of energy consumption per bird type

A second expression for a bird type’s energy consumption at a given location during a prescribed period is the product of the number of birds times the mean energy intake per bird, as expressed by the following formula:

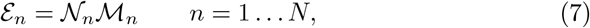

where 𝒩_*n*_ and ℳ_*n*_ represent respectively the total number and mean energy consumption of birds of type *n*.

As with transect observations, we assume that point counts are representative samples across all bird types, so we have:

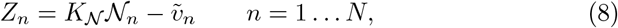

where

- *K*_𝒩_ : Constant of proportionality, assumed to be independent of bird types;
- *Z*_*n*_: Number of point count observations of bird type *n, n* = 1 … *N* ;
- 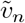: Uncertainty in point counts for bird type *n*.

As with transect observations, we assume that the uncertainty is proportional to the number of observations:

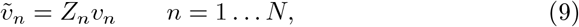

where {*v*_*n*_, *n* = 1 … *N* } are independent, identically distributed random variables with mean 0 and variance 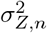.

In the literature, bird type mean energy consumption is modeled as proportional to mean bird weight raised to a power as follows [14]:

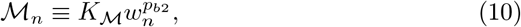

where

- *K*_ℳ_: Constant of proportionality, independent of bird type *n*;
- *w*_*n*_: Mean weight of bird type *n, n* = 1, … *N* ;
- *p*_*b*2_: Weight exponent in energy model.

By substituting (8)-(10) into (7) we obtain finally:

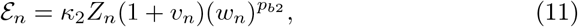

where *κ*_2_ ≡ *K*_ℳ_*/K*_𝒩_ is independent of *n*.

#### 3.1.3 Parameter estimation based on energy consumption equations

In practice, equations (5) and (11) developed in the previous two sections cannot be used to estimate total energy consumption of bird types, because the constants *κ*_1_ and *κ*_2_ are impossible to obtain. However, by setting the two expressions equal to each other we obtain a relationship that enables us to estimate the exponents *p*_*b*1_, *p*_*b*2_ and *p*_*f*_ based on observed point count and transect data. Equating ℰ_*n*_ in (5) and (11) gives:

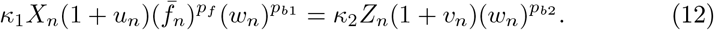

Taking logarithms and rearranging, we find:

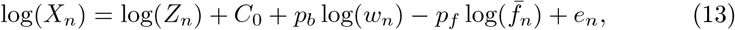

where:

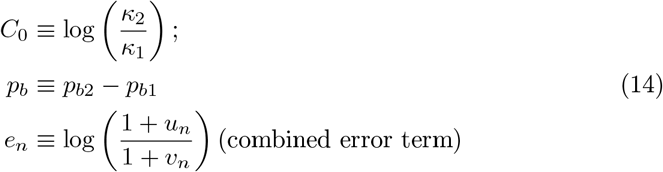

Note that if *u*_*n*_, *v*_*n*_ ≪ 1 then *e*_*n*_ ≈ *u*_*n*_ − *v*_*n*_. We will assume that {*e*_*n*_, *n* = 1, …*N* } are independent, identically distributed random errors. Then if we have transect data {*X*_1_, …, *X*_*N*_ } and point count data {*Z*_1_, …, *Z*_*N*_ }, we may use (13) to perform ordinary unweighted least squares regression and find values for *C*_0_, *p*_*b*_, and *p*_*f*_ that give the best fit to the data.

#### 3.1.4 Estimates of relative abundance

In this section we make use of the results from previous sections to develop a pair of estimates for relative abundances of bird types, where the relative abundance *a*_*n*_ of species *n* is defined as type abundance divided by total bird abundance:

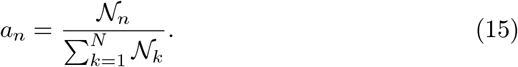

The two estimates are based on point counts and transect observations, respectively. We also show that in the case where the estimators are unbiased and uncorrelated, a linear combination of the two estimators can be formed that is unbiased and has smaller variance than either of the two individual estimators.

First we develop an estimator for relative abundance based on point counts.

From (8) and (9) and the definition (15) we have:

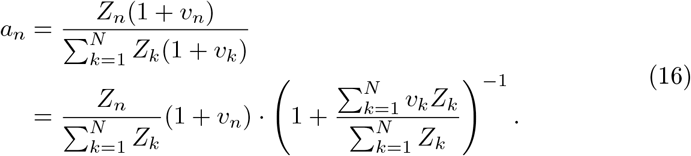

The final multiplicative term in (16) is independent of *n*, and involves the sum of independent random errors, which will tend to cancel each other. We may therefore approximate this term as:

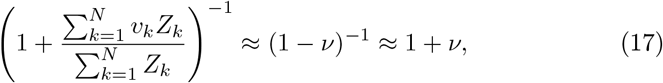

where *ν* ≪ 1 is a mean-zero random variable. This gives finally

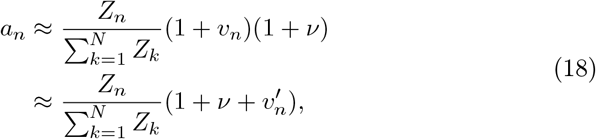

where 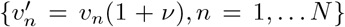 are statistically independent errors. It follows that an estimator 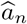 for *a*_*n*_ that is based on point counts {*Z*_*n*_} can be obtained from (18) with the random error terms removed:

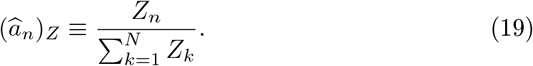

Next, an alternative expression for *a*_*n*_ based on transect observations can be obtained. Combining (8), (9) and rearranging, we have:

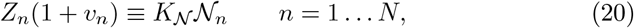

and substituting into (12) and rearranging gives:

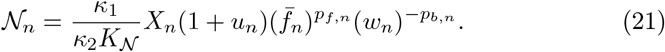

(Note that we are generalizing (12) to the case where the fruit weight and bird weight exponents *p*_*f,n*_ and *p*_*b,n*_ depend on bird type.) This enables us to give an expression for relative abundance in terms of *X*_*n*_ instead of *Z*_*n*_:

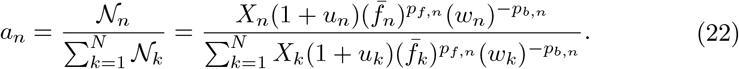

By a similar argument as before, we obtain the following *X*_*n*_-based estimator for relative abundance *a*_*n*_:

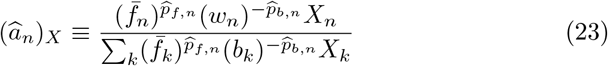

where 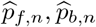 are estimates for the fruit and bird weight coefficients *p* _*f,n*_ and *p*_*b,n*_, respectively.

#### 3.1.5 Minimum variance combination of relative abundance estimators

Now consider the linear combination of estimators:

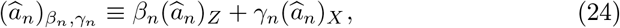

where {*β*_*k*_, *γ*_*k*_, *k* = 1 … *N* } are positive coefficients. It follows by the linearity of distribution mean that:

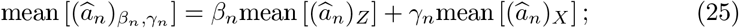

In the case where the estimators 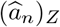 and 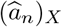 are unbiased, we have:

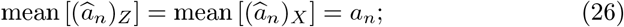

The requirement that 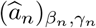 be an unbiased estimator of *a*_*n*_ is expressed mathematically as:

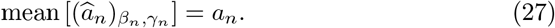

Combining (25), (26), and (27) gives the requirement:

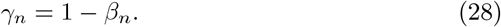

Given that 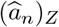 and 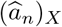 are based on independent observations and are thus uncorrelated, it follows:

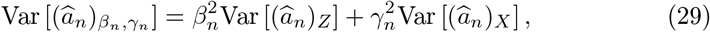

and a standard calculation shows that the unbiased minimum-variance estimator for *a*_*n*_ is obtained by choosing:

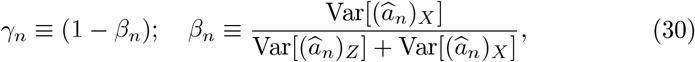

which has variance

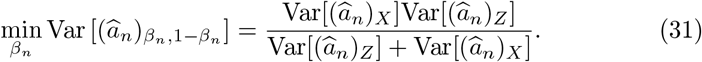

#### 3.1.6 Combined estimates of log relative abundance

Sometimes it is more advantageous to consider the logarithms of relative abundances of bird species, genera, or families, instead of relative abundances themselves. Typically the logarithms have a more symmetrical distribution which is closer to normal. By taking natural logarithms of 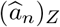 and 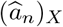, we obtain estimators for the logarithm of relative abundance, which we denote as 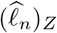 and 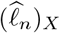 respectively:

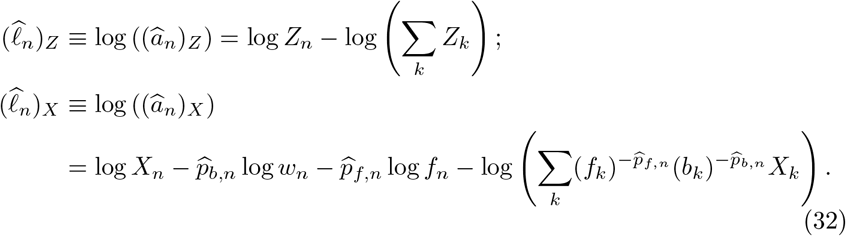

As with the relative abundance estimators, we may form linear combinations of the log relative abundance estimators:

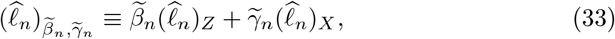

where 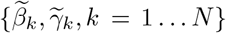 are positive coefficients. In the case where 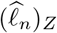 and 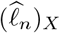 are unbiased and uncorrelated, it follows as before that:

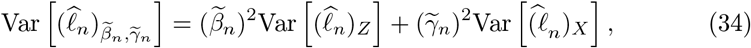

and an unbiased minimum-variance estimator for *ℓ*_*n*_ is obtained by choosing:

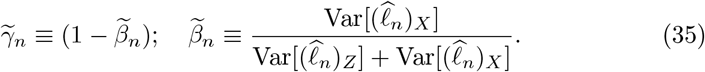

#### 3.1.7 Calculation of coefficients for combined estimators

The combined estimators of relative abundance and log relative abundance defined in Sections 3.1.5 and 3.1.6 respectively require the specification of coefficients 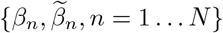. These coefficients are not determined a priori, but rather must be supplied by the user. In this section, we describe a simulation-based procedure for determining coefficients which will give estimators that have reduced variance.

The proposed estimation procedure for *β*_*n*_ is based on a bootstrapping approach, and runs as follows. Suppose the user has recorded point counts 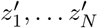 and transect observations 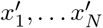 for *N* different species.

1. Compute abundance estimates based on 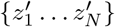, according to (19):

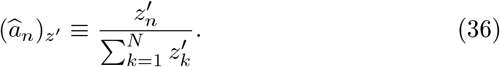
2. Generate *M* simulated sets of point counts and transect observations 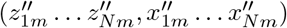, *m* = 1 … *M* based on the user’s observed data. Simulation values are independently generated according to the following distributions:

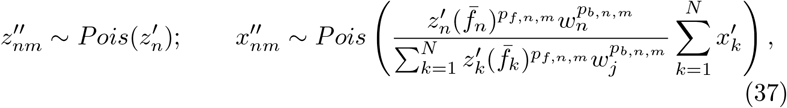

where *Pois*(*λ*) denotes a Poisson random variable with mean event rate *λ*; and *p*^*′*^_*f,n,m*_ and *p*^*′*^_*b,n,m*_ are randomly generated according to probability distributions reflecting the user’s knowledge of fruit and weight exponents in the model (3). (Note that Poisson distributions are commonly used to model survey counts [15].) This expression generates simulated point counts with expected values equal to the observed point counts 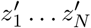, and simulated transect observations whose expected values yield energy consumption rates that agree with those calculated from the point counts, and whose sum agrees with the sum of the user’s observed transect observations.
3. For each of the *M* simulated sets of observations, the user calculates abundance estimators according to (19):

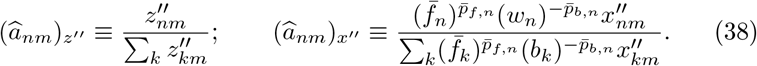
4. The user calculates values of *β*_*n*_, *n* = 1 … *N* that minimize root mean squared error (RMSE) of the combined estimator 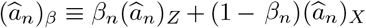:

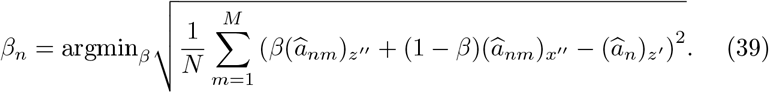

The minimized RMSE values in (39) can also be used to estimate confidence intervals for 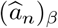, since the RMSE value should approximate one standard deviation. So if the RMSE value obtained is denoted as Δ_*β,n*_, then 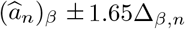 should give a 90% confidence interval for the true value of *a* .

A similar procedure can be used to determine values of 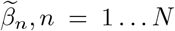 and confidence intervals for an optimal linear combination of log abundance estimators, defined as 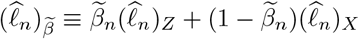.

### 3.2 Empirical evaluation and verification of theoretical equations

In this section, we describe how the equations derived in Section 3.1 are evaluated and verified using data from an actual survey of frugivore birds conducted in several locations in Ecuador. Both the data collected and the verification methods applied are described.

#### 3.2.1 Dataset used in methodology evaluation

Data was collected from 18 one-hectare plots spread across three elevations: approximately 1000 m above sea level (asl) (970–1092 ± 47.2 m, 4^°^6^*′*^S, 78^°^58^*′*^W); 2000 m asl (2025–2119 ± 32.4 m, 3^°^58^*′*^S, 79^°^4^*′*^W); and 3000 m asl (2874–2966 ± 32.9 m, 4^°^6^*′*^S, 79^°^10^*′*^W). These plots represented two types of habitat—natural forest and fragmented forest—within and around Podocarpus National Park (PNP) and the San Francisco Biological Reserve (BRSF) on the southeastern slope of the Andes in southern Ecuador. Podocarpus National Park is recognized as one of Ecuador’s most biodiverse protected areas and encompasses a steep elevational gradient supporting distinct vegetation zones, from evergreen premontane forest at low elevations through evergreen lower montane forest at mid elevations to upper montane forest dominated by *Weinmannia* shrubs, vascular epiphytes, and epiphytic mosses at high elevations [16, 17]. The fragmented sites were located in working landscapes surrounding the park, where the primary land use separating forest patches consists of cattle pasture and small-scale agriculture, creating a mosaic of remnant forest fragments embedded in open matrix habitat. The six sites are characterized in Table 1. Study plots were selected jointly by the German Platform for Biodiversity and Ecosystem Monitoring and Research in South Ecuador (RESPECT) to ensure representative coverage of each forest type across the elevational gradient. The area exhibits a humid tropical montane climate [18] with a bimodal rainfall pattern, with peak precipitation from May through June and minimum precipitation from October through November [19].

**Table 1:** Summary of characteristics for the six study locations. Natural sites are located within protected forest; fragmented sites are located in working landscapes where remnant forest patches are separated by cattle pasture and small-scale agriculture.

| Location | Approx. altitude (m) | Environment type |
| --- | --- | --- |
| Bombuscaro | 1000 | Natural |
| Copalinga | 1000 | Fragmented |
| ECSF | 2000 | Natural |
| Finca | 2000 | Fragmented |
| Cajanuma | 3000 | Natural |
| Bellavista | 3000 | Fragmented |

##### Point count surveys

To assess bird diversity and abundance, nine observation points were established within each one-hectare plot, spaced at least 50 m apart to minimize double-counting. At each point, a trained observer recorded all bird species detected by sight or sound within a 20-m radius during a fixed 10-minute observation window [20]. Surveys were conducted during the first three hours after sunrise, when bird activity is highest, and were suspended during rain or strong wind. Each plot was surveyed eight times over a two-year period (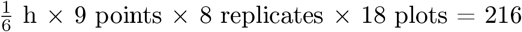 total sampling hours). To minimize observer bias, all surveys at a given plot were conducted by the same trained observer. The total number of bird species (species richness) and overall number of individual birds (abundance) at each plot were quantified by aggregating records across all observation points and replicates. The point count dataset comprises a total of [*N*_pc_] individual detection records across all plots and replicates. Bird species were classified as frugivores if their diet consisted of 40% or more fruit [21].

##### Transect surveys

Transect surveys were conducted along fixed linear transects of 200 m established within each plot. Observers walked each transect at a slow, steady pace (approximately 1 km/h) during the same morning activity window used for point counts. All frugivore feeding events observed within 10 m of the transect centerline were recorded, including the identity of the bird (to species level where possible, otherwise to genus or family), the plant species being consumed, and the number of fruit items ingested per feeding bout. Each transect was surveyed on the same eight occasions as the point count plots. The transect dataset comprises a total of [*N*_tr_] feeding event records. Mean body masses for each bird species or family were obtained from published sources [21], and mean fruit masses for each plant species consumed were obtained from field measurements or the literature.

#### 3.2.2 Parameter estimation using energy models

Best fits for the parameters *p*_*f*_ and *p*_*b*_ were found by applying linear regression to (13), for several different datasets. Fits were found for species data from each of the six regions separately; for species data grouped by altitude; for species data grouped by environment type (natural versus fragmented); and for species data for all six regions combined. For each fit, scatter plots were made for actual versus predicted transect observations, based on (13). *R*^2^ and Akaike information criterion (AIC) values and 95 percent confidence intervals for *p*_*f*_ and *p*_*b*_ were also computed. We use AIC rather than likelihood-ratio tests or *p*-values for model comparison because AIC penalizes model complexity in a way that is directly interpretable in terms of information loss, and because the one- and two-parameter models compared here are not strictly nested in a manner that satisfies the assumptions of a formal likelihood-ratio test; a smaller AIC value indicates that the added parameter does not justify the increased model complexity [22].

Scatter plots, *R*^2^ and AIC values and 95 percent confidence intervals for *p*_*b*_ were also generated for the same datasets, but with a simplified single-parameter model that assumes *p*_*f*_ = 0 in (13). The 2-parameter and 1-parameter models are compared for goodness of fit. The purpose is to determine whether *p*_*f*_ has a significant effect on mean energy consumption per feeding. All computations were performed in Python (version 3.9) using the `NumPy`, `SciPy`, and `statsmodels` libraries. Simulation code implementing the bootstrapping procedure described in Section 3.1.7 is available at [repository URL].

#### 3.2.3 Performance evaluation of abundance estimation using empirically-based simulations

Statistical characteristics of the abundance and log abundance estimators defined in Sections 3.1.4 and 3.1.6 were evaluated using simulations as follows. In this evaluation, the purpose was not to estimate true relative abundances of the regions from which the dataset was taken—since we don’t know these true relative abundances, the accuracy of such estimates cannot be ascertained. Rather, the regional data is used to supply realistic scenarios; the regional data from the six regions is used as “ground truth” for six hypothetical regions, and simulations are randomly generated based on the data, and used to predict this “ground truth”. In this way we can evaluate the accuracy of estimators using simulations with realistic characteristics. We base simulations on per-family data instead of per-species data, because per-species point counts are too small to establish differences in accuracies between estimators.

##### Testing for estimator accuracy and precision

Simulations were conducted to determine the accuracy and precision of point count-based, transect observation-based, and combined estimators as follows.

10000 simulated scenarios were generated based on the bird family data from each region. The data from the six actual regions was used to generate “ground truth” for six hypothetical regions as follows. If *PC*_1_, *PC*_2_, … *PC*_*N*_ are the observed point counts in a given actual region, then the true relative abundance of family *n* in the corresponding hypothetical region is given by:

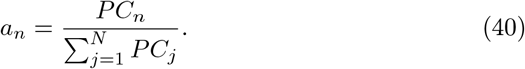

Next, we simulated 10,000 surveys in each hypothetical region, where each survey includes point counts and transect observations. For a given simulation, the point count for each bird family *n* is generated by a Poisson random variable whose mean is the point count from the observed data. We denote these Poisson random variables as *z*_*n*_, *n* = 1, … *N*, where

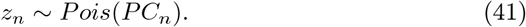

Transect observations for each simulation are randomly generated as follows. First, for each bird family *n* we generate a weight exponent *p*_*b,n*_ as a Gaussian random variable with 95% confidence interval [−0.6, −0.1]:

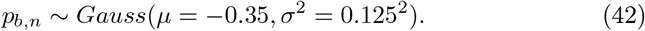

The choice of distribution (42) will be motivated in Section 4.1 (see the final line of Table 3): this represents the uncertainty in the parameter *p*_*b*_ which is estimated as described in Section 3.2.2. The transect observation for birds in family *n* (denoted as *x*_*n*_) is then simulated as follows:

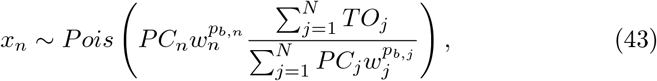

where *TO*_*j*_ denotes the number of transect observations of bird family *j* in the original data (*j* = 1 … *N*). The choice of Poisson parameter in (43) was made so that the expected sum of simulated observations over bird families at a given region is equal to the sum of transect observations in the original data. The Poisson parameter also reflects a 1-parameter model of energy consumption that presumes *p*_*f*_ = 0 in (4). This choice will be justified by the results in Section 4.1, which show there is no statistical evidence of energy consumption’s dependence on mean fruit weight.

Given these simulated point counts {*z*_*n*_} and transect observations {*x*_*n*_}, we then compute estimates 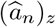 and 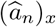 for *a*_*n*_ based on {*z*_*n*_} and {*x*_*n*_} respectively according to (19) and (23):

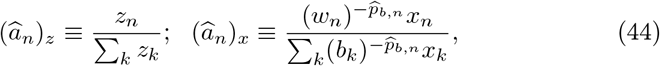

where 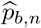 is the investigator’s best determination of the bird weight exponent *p*_*b,n*_. Two cases were considered: (a) the investigator does not have detailed knowledge of the bird weight exponent, and sets 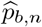 equal to the mean value 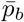 of the distribution in (42); and (b) the investigator knows exactly the values of *p* _*b,n*_ for each *n*, and uses 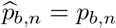.

Besides the estimators 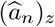 and 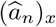, we also computed the combined estimate 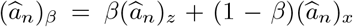 as a function of *β* for 0 ≤ *β* ≤ 1 for each simulation. Taking these results over all simulations, we were able to compute the means and variances of 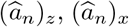, and 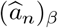. This enables us to compute the accuracy and precision of all estimators for each family in each of the six hypothetical regions, for the two cases (a) and (b) defined in the previous paragraph.

A similar procedure was used to gauge the accuracy and precision of log abundance estimators 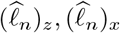, and 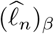 defined as follows:

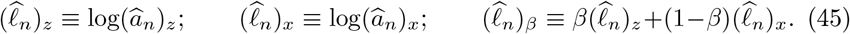

##### Combined estimator parameter determination and confidence interval estimation

The second set of simulations is intended to test the procedure for determining variance-reducing linear combinations of estimators described in Section 3.1.7.

- First, for each of the six different regions observed in the study, 2500 simulated scenarios for point count and transect observation values {*z*_1_ … *z*_*n*_, *x*_1_, … *x*_*n*_ }_*m*_, *m* = 1 … 2500 were generated according to (41). Each observation vector *z*_1_ … *z*_*N*_ was used to create relative abundance vectors 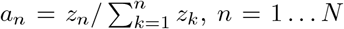. These relative abundances are taken as “ground truth” for the scenario (these are generated so as to increase the variety in the “ground truth” scenarios, making the test more robust.)
- Next, for each scenario, point counts and transect observations (simulating the dataset collected by a single hypothetical “ecologist” studying the scenario) were randomly generated as 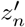 and 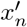 respectively according to (compare (41)-(43))

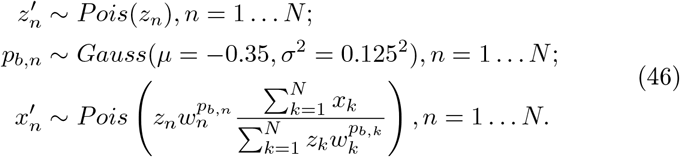
- Each simulated ecologist uses her own point counts and transect observations to compute estimators 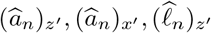, and 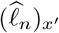 according to (38). Two alternative situations are modeled: (a) the ecologist does not have detailed knowledge of the coefficients *p*_*b,n*_, and uses the mean value 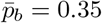 to estimate 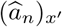 and 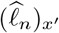; (b) the ecologist has precise knowledge of *p*_*b,n*_, and uses the same values that are used in (46) for these estimates.
- Each simulated ecologist then uses the prescription in Section 3.1.7 with *M* = 2500 to estimate the linear combination parameters *β* and 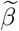 for the estimators 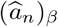 and 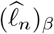, as well as 90% confidence intervals for these estimates. As in the previous step, two scenarios apply: (a) the ecologist uses the mean value 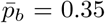 to estimate 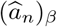 and 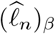 (b) the ecologist uses the same values of *p*_*b,n*_ that are used in (46) for these estimates.

To compare the accuracies of the point count-based, transect-based, and combined estimators, 95% confidence intervals for the relative error in relative abundance (estimate − ground truth relative abundance)/(relative abundance) are plotted. In order to verify the confidence intervals computed in the last bullet above, we also recorded the proportion of time that the ground truth relative abundances were within the 90% confidence intervals.

## 4 Results

### 4.1 Application of energy consumption estimation methodology to dataset

The six panels in Figure 1 show 95% confidence intervals (± 2 standard deviations) for the model coefficients *p*_*b*_ (*left*) and *p*_*f*_ (*right*) as estimated from different data subsets using the two-coefficient model (13). Confidence intervals are shown for the coefficient estimates using per-species data from six separate locations (yellow bars); using data grouped by altitude (purple bars); using data grouped by fragmentation (blue bars); and using all data combined (green bar). In 11 of the 12 cases (including the overall data model), the confidence intervals for *p*_*f*_ contain 0, indicating a lack of statistical evidence for a nonzero value of *p*_*f*_ . Equivalently, this result is consistent with a one-sided test of *H*_0_ : *p*_*f*_ = 0 at the 5% significance level in 11 of 12 cases. On the other hand, in 6 of the 12 cases (including the model for overall data), the confidence interval for *p*_*b*_ lies entirely below 0, indicating that *p*_*b*_ is negative at 5% significance.

**Figure 1:**
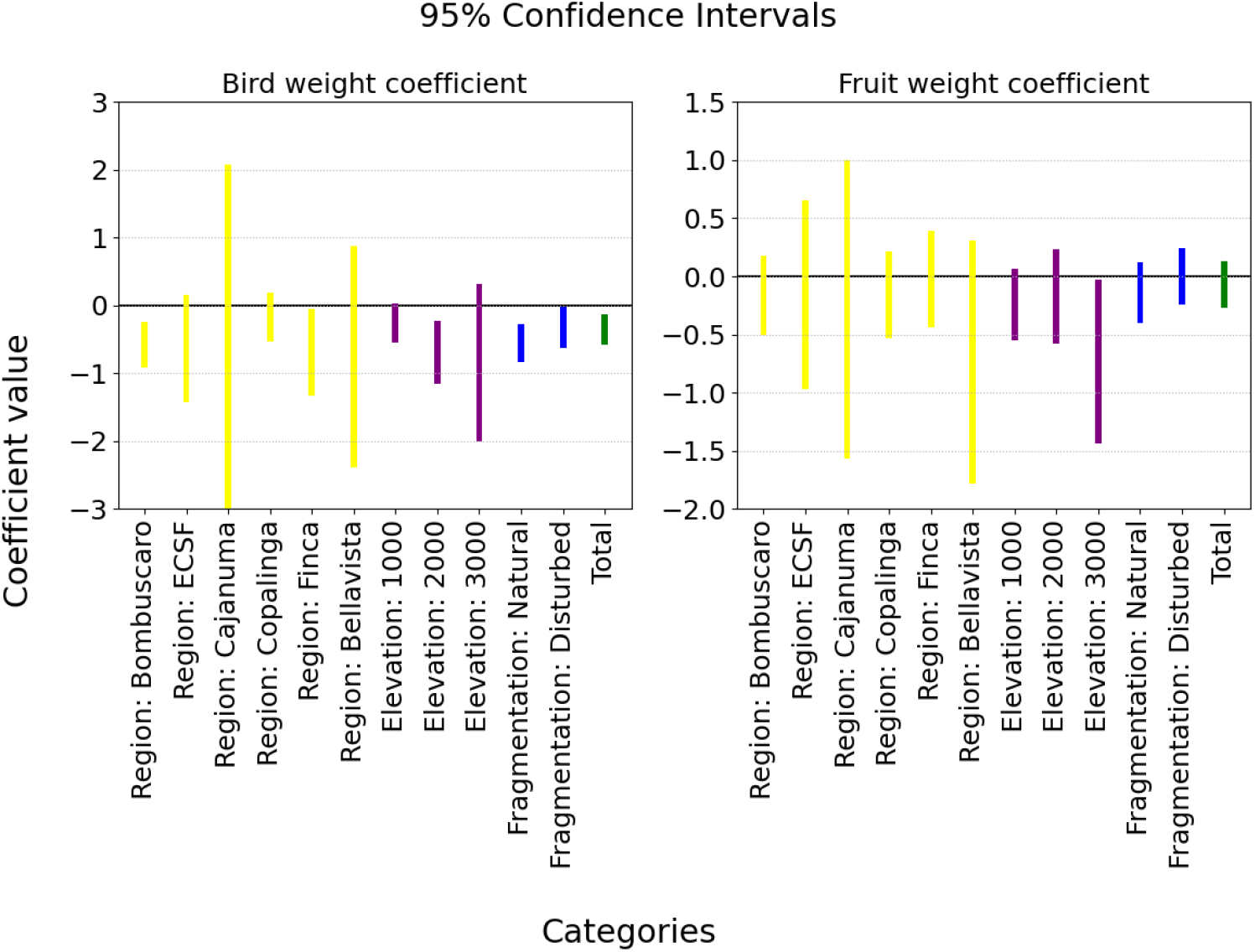
95% confidence intervals for the bird weight coefficient *p*_*b*_ (*left panel*) and the fruit weight coefficient *p*_*f*_ (*right panel*) from the two-coefficient model (13), estimated from different data subsets: by site (yellow bars, *n* = 6), by elevation (purple bars, *n* = 3), by habitat type (blue bars, *n* = 2), and for all data combined (green bar, *n* = 1). Confidence intervals for *p*_*f*_ that contain zero indicate no statistically significant effect of fruit weight on feeding frequency at *α* = 0.05; this was the case in 11 of 12 subsets. Confidence intervals for *p*_*b*_ lying entirely below zero indicate a significant negative effect of bird weight, consistent with larger birds consuming more energy per feeding event.

Scatter plots of actual versus predicted transect observations for the two-coefficient model, by location, elevation, habitat type, and combined, are provided in Supplement A (Figures A1–A4). For most regions there is a generally positive trend, albeit with numerous errant values particularly on the low side, and there is no discernible influence of bird weight on the accuracy of the predictions. A summary of the best-fit parameters, *R*^2^ values, and AIC values for the two-coefficient model across all data subsets is given in Table 2. *R*^2^ values range from 0.03 to 0.40, indicating relatively poor fits.

**Table 2:**
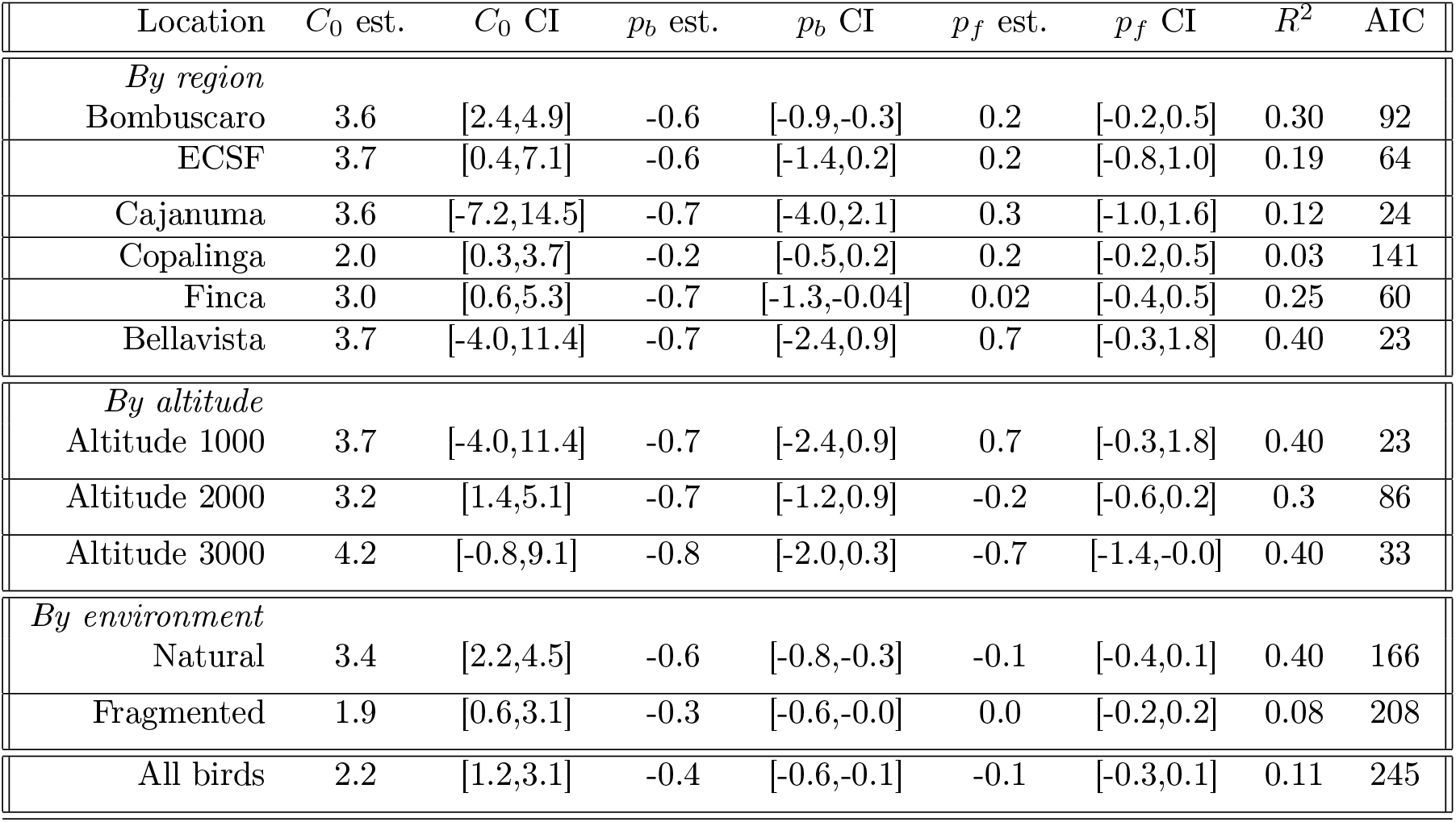
Summary of parameter estimations from regression for the two-coefficient model (13), which includes both bird weight and fruit weight. CI = 95% confidence interval; AIC = Akaike information criterion.

**Table 3:** Summary of parameter estimations from regression for the single-coefficient model with *p*_*f*_ = 0 in (13). CI = 95% confidence interval; AIC = Akaike information criterion. Reduced AIC values relative to Table 2 confirm that dropping the fruit weight parameter improves model parsimony.

| Location | $C_0$ est. | $C_0$ CI | $p_b$ est. | $p_b$ CI | $R^2$ | AIC |
| --- | --- | --- | --- | --- | --- | --- |
| *By region* |  |  |  |  |  |  |
| Bombuscaro | 3.4 | [2.2,4.5] | -0.6 | [-0.9,-0.2] | 0.27 | 91 |
| ECSF | 3.4 | [0.4,7.1] | -0.6 | [-1.3,0.2] | 0.18 | 63 |
| Cajanuma | 1.7 | [-3.9,7.3] | -0.3 | [-1.9,1.4] | 0.03 | 22 |
| Copalinga | 1.5 | [0.28,2.7] | -0.14 | [-0.5,0.2] | 0.01 | 140 |
| Finca | 2.9 | [0.8,5.0] | -0.7 | [-1.3,-0.0] | 0.25 | 58 |
| Bellavista | -0.9 | [-5.4,3.6] | 0.09 | [-1.15,1.33] | 0.00 | 25 |
| *By altitude* |  |  |  |  |  |  |
| Altitude 1000 | 1.9 | [0.9,2.9] | -0.3 | [-0.5,0.0] | 0.06 | 156 |
| Altitude 2000 | 2.9 | [1.2,4.6] | -0.6 | [-1.1,-0.2] | 0.27 | 85 |
| Altitude 3000 | 0.2 | [-3.7,4.1] | -0.1 | [-1.1,1.0] | 0.00 | 37 |
| *By environment type* |  |  |  |  |  |  |
| Natural | 3.1 | [2.1,4.0] | -0.5 | [-0.8,-0.25] | 0.22 | 166 |
| Fragmented | 1.9 | [0.9,2.9] | -0.3 | [-0.6,0.0] | 0.08 | 206 |
| All | 2.0 | [1.2,2.8] | -0.3 | [-0.6,-0.1] | 0.1 | 244 |

The fact that *p*_*f*_ is not significantly different from 0 in virtually all cases motivates replacing *p*_*f*_ = 0 in (13). The model then has only the coefficient *p*_*b*_ plus the constant *C*_0_ that are fitted to the transect and point count data:

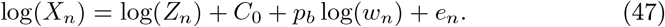

Figure 2 summarizes confidence intervals for the bird weight coefficient in the single-coefficient model (47). Confidence intervals are consistent across all regions, and overlap in the range of −0.3 to − 0.5. As with the two-coefficient model, the bird weight coefficients tend to be negative, and some are negative with 95% significance (including the overall estimate). Confidence intervals for high altitudes are smaller, which may reflect the fact that bird species are more variable at low altitudes. Taken together, the data gives a 95% confidence interval for *p*_*b*_ as [−0.6, −0.1]. Scatter plots for the one-coefficient model, by location, elevation, habitat type, and combined, are provided in Supplement A (Figures A5–A8). Visually, the one-coefficient model appears to have accuracies comparable to the two-coefficient model.

**Figure 2:**
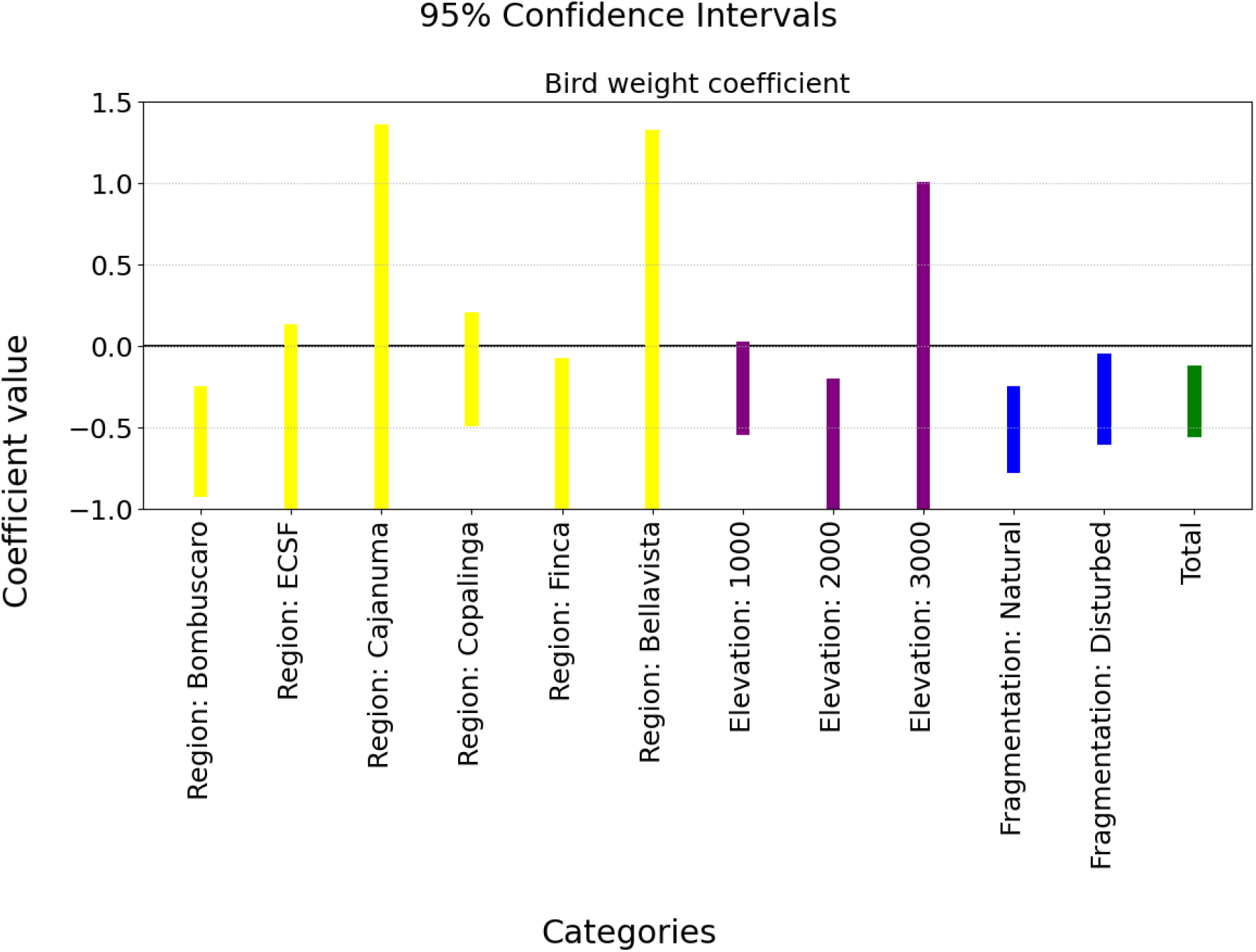
95% confidence intervals for the bird weight coefficient *p*_*b*_ in the one-coefficient model (47) (*p*_*f*_ = 0), estimated from different data subsets: by site (yellow), by elevation (purple), by habitat type (blue), and for all data combined (green). Confidence intervals are broadly consistent across subsets, overlapping in the range [−0.3, −0.5]. The combined dataset yields a 95% confidence interval of [−0.6, −0.1], used to parameterize the simulation distributions in Section 3.2.3.

Table 3 summarizes the fit parameters for the single-coefficient model. The AIC values for all fits are reduced relative to the two-coefficient model (Table 2), thus confirming that the exclusion of fruit weight as a significant parameter is justified: a smaller AIC indicates that the loss of information from dropping *p*_*f*_ is more than offset by the reduction in model complexity.

Recall that *p*_*b*_ is defined in (14) as the difference of the two exponents *p*_*b*1_ and *p*_*b*2_, where *p*_*b*1_ is related to bird weight’s influence on energy consumption per bird, while *p*_*b*2_ is related to bird weight’s influence on energy consumption per observed feeding. Estimates of *p*_*b*1_ in [14] for most families are close to 0.7 (hummingbirds are an exception, with *p*_*b*1_ *>* 1). Our finding of *p*_*b*_ ≈ − 0.4 is consistent with a value of *p*_*b*1_ ≈ 1. This would imply that bird energy consumption per feeding is roughly proportional to bird weight. This is not unreasonable, because heavier birds proportionately have larger anatomical features, including larger stomachs.

Figure 3 shows a scatter plot of the ratio of feeding observations to point count observations versus point count frequency. The figure indicates a negative correlation between the feeding/observation ratio and the number of observations. This is an indication of regression to the mean: overpredictions (or under-predictions) based on transect observations will tend to be revised downwards (or upwards) based on feeding observations—and vice versa. The presence of regression to the mean indicates that point counts and transect observations tend to counterbalance each other, and estimators based on both factors may achieve greater accuracy. (This indication is confirmed in the following sections.)

**Figure 3:**
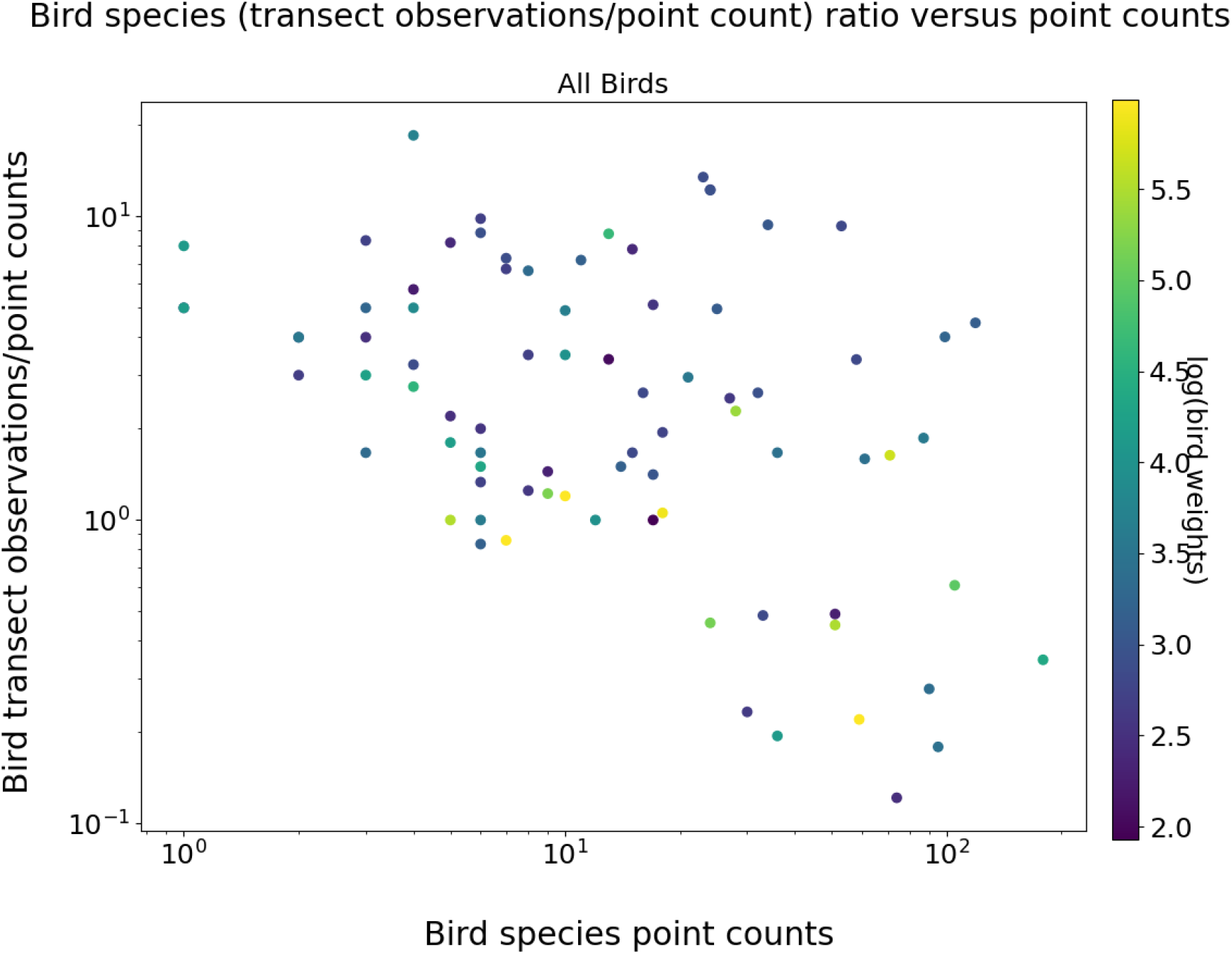
Scatter plot of the per-family ratio of feeding observations to point count observations versus total point count observations, using data pooled across all six sites. Each point represents one bird family at one site. The negative trend indicates regression to the mean: families with high point counts tend to have proportionally fewer feeding observations, and vice versa, suggesting the two data types provide complementary and partially counterbalancing information about abundance.

### 4.2 Performance evaluation of estimators

The results of the performance evaluation tests described in Section 3.2.3 are summarized in Figures 4 and 5–6 in the main body, with full supporting detail provided in Supplements B and C. Supplement B contains results for estimators computed using the mean value 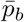 of the bird weight exponent (case a), while Supplement C contains corresponding results using the exact values of *p*_*b,n*_ (case b), as defined in Section 3.2.3.

**Figure 4:**
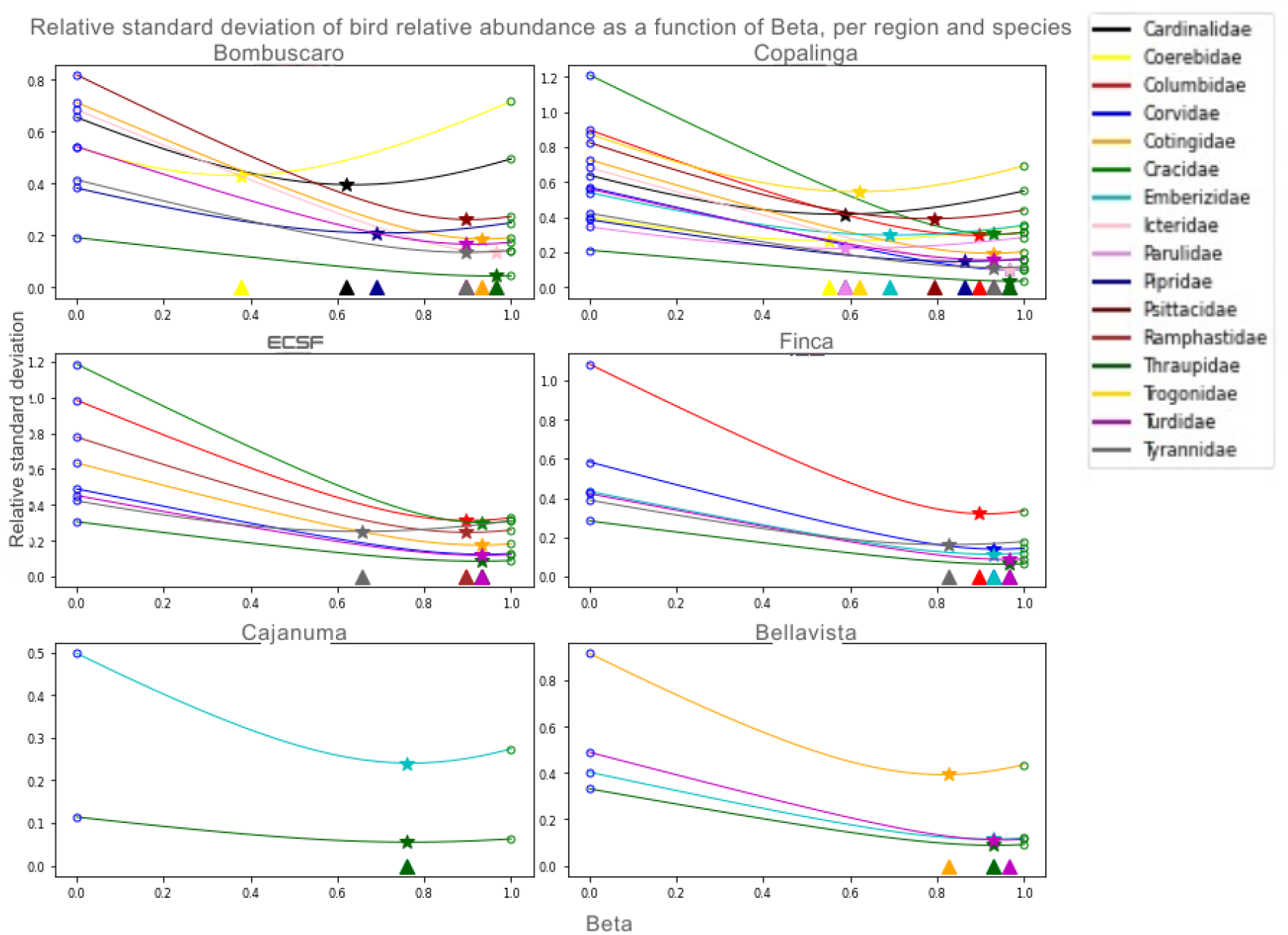
Relative standard deviation (standard deviation divided by true relative abundance) of the combined estimator 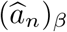 as a function of the mixing parameter *β* ∈ [0, 1], for each bird family in each of the six study sites (one curve per family). *β* = 0 corresponds to the pure transect-based estimator 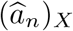; *β* = 1 corresponds to the pure point count-based estimator 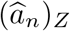. Stars mark the minimum of each curve; triangles on the *x*-axis indicate the theoretically optimal *β* from (30). The transect estimator uses the mean bird weight exponent 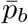 (case a). The close agreement between stars and triangles confirms the accuracy of the analytical formula.

**Figure 5:**
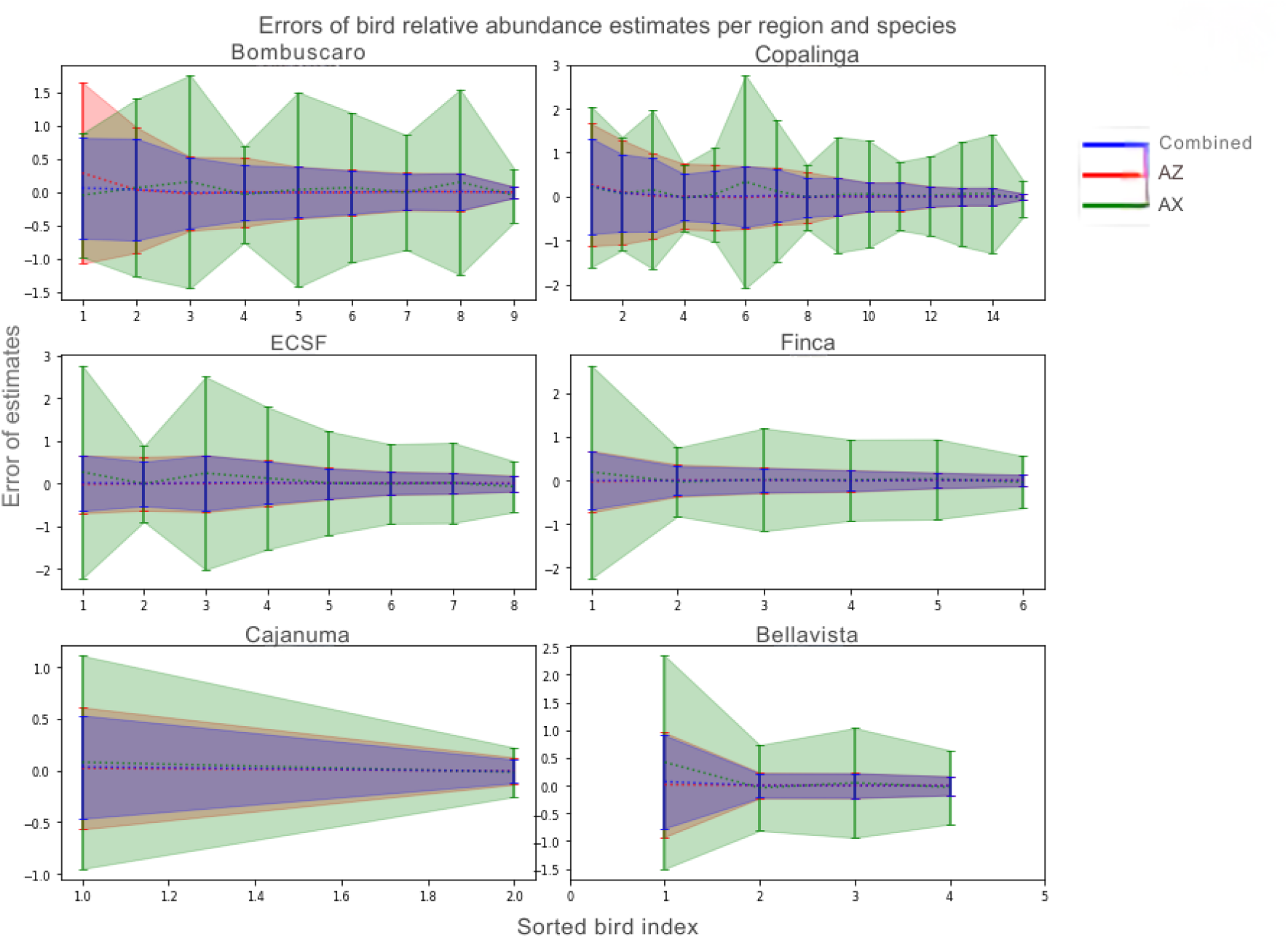
95% confidence intervals for the relative error (estimated minus true relative abundance, divided by true relative abundance) of three estimators: the point count-based estimator 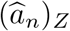 (labeled *AZ*), the transect-based estimator 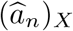 (labeled *AX*), and the combined estimator 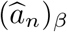. Results are shown for each bird family in each of the six study sites. The *AX* and combined estimators use the mean bird weight exponent 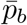 (case a). Families are ordered by increasing true relative abundance within each site panel. The combined estimator confidence interval is always at least as narrow as the better of the two individual estimators.

#### Case (a): estimators computed using mean value of bird weight exponent 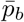

The mean values of the estimators 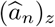 and 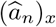 over 10,000 simulations, compared to the true relative abundance, are shown in Figure B1 of Supplement B. The figure shows that there is very little bias in most estimators, with estimators for birds with lower relative abundance tending to have the most bias. Figure B2 of Supplement B shows similar results for the log abundance estimators 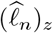 and 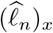.

Figure 4 shows the relative standard deviations for 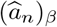 for values of *β* between 0 and 1, for all different bird families in the six different regions. The standard deviations are based on estimates of 10,000 simulated ecologists. The triangles on the horizontal axis mark the optimal values of *β* as calculated by (30). The figure shows that (30) gives a very accurate estimate of the *β* that minimizes standard deviation. All of the optimal *β* values are between 0.5 and 1, reflecting the fact that 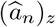 (corresponding to *β* = 1) is consistently more accurate than 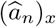 (corresponding to *β* = 0). Figure B3 of Supplement B shows similar results for the standard deviations for 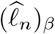.

Figures B4 and B5 of Supplement B show the relative RMSE of 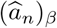 and 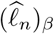 respectively as functions of *β*. These graphs are nearly identical to the corresponding standard deviation figures, confirming that bias error is negligible compared to standard deviation for both estimators.

Figures B6 and B7 of Supplement B show RMSE-minimizing *β* values for 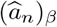 and 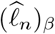 respectively, versus the true relative abundance of each bird family. It is clear that lower *β* values (between 0.4 and 0.9) only occur for birds with relative abundance less than about 0.1, with only two exceptions from the Cajanuma dataset (which had only 2 families).

#### Case (b): Estimators computed using exact values of bird weight exponent p_b,n_

Supplement C contains figures corresponding to those in Supplement B, except they show results obtained when the investigator has accurate information about *p*_*b,n*_. It is evident from Figures C3–C6 of Supplement C that the estimates 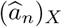 and 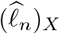 (corresponding to *β* = 0) are more accurate than the corresponding estimates when *p*_*b,n*_ is not known exactly. As a result the optimal values of *β* tend to be lower when *p*_*b,n*_ is known exactly, as shown in Figures C7 and C8 of Supplement C (compare Figures B6 and B7 of Supplement B for the inexact case).

### 4.3 Verification of accuracy of combined estimators

Figures 5 and 6 show 95% confidence intervals for the relative errors for point count-based, transect observation-based, and combined estimators of relative abundance obtained from the simulations described in Section 3.1.7; results are also summarized numerically in Tables 4 and 5. Both figures show that relative abundance estimates using the combined estimator 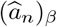 are always at least as accurate as the best of the two estimators 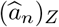 and 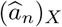. Comparing between the two figures shows that combined estimates using the exact values for *p*_*b,n*_ are much more accurate than those using 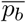.

**Table 4:** Means and standard deviations of RMSE for relative error of point count-based, transect-based, and combined relative abundance estimators, for individual regions and grouped by altitude, environment type, and total.

| | $\hat{a}_Z$ rel. err. | | $\hat{a}_X$ rel. err. | | $\hat{a}_\beta$ rel. err. | |
| --- | --- | --- | --- | --- | --- | --- |
| | (mean ± std) | | mean $\bar{p}_b$ | exact $p_b$ | mean $\bar{p}_b$ | exact $p_b$ |
| Bombuscaro | 0.28 ± 0.21 | | 0.55 ± 0.18 | 0.16 ± 0.11 | 0.22 ± 0.12 | 0.14 ± 0.10 |
| Copalinga | 0.30 ± 0.19 | | 0.64 ± 0.27 | 0.23 ± 0.17 | 0.25 ± 0.15 | 0.18 ± 0.13 |
| ECSF | 0.23 ± 0.10 | | 0.66 ± 0.29 | 0.21 ± 0.12 | 0.21 ± 0.15 | 0.15 ± 0.08 |
| Finca | 0.16 ± 0.10 | | 0.54 ± 0.28 | 0.16 ± 0.14 | 0.15 ± 0.09 | 0.11 ± 0.08 |
| Cajanuma | 0.18 ± 0.11 | | 0.33 ± 0.21 | 0.14 ± 0.09 | 0.16 ± 0.10 | 0.10 ± 0.07 |
| Bellavista | 0.20 ± 0.16 | | 0.60 ± 0.34 | 0.49 ± 0.50 | 0.20 ± 0.16 | 0.18 ± 0.15 |
| Altitude 1 | 0.29 ± 0.20 | | 0.61 ± 0.24 | 0.20 ± 0.15 | 0.24 ± 0.14 | 0.16 ± 0.12 |
| Altitude 2 | 0.20 ± 0.10 | | 0.61 ± 0.29 | 0.19 ± 0.13 | 0.18 ± 0.09 | 0.13 ± 0.08 |
| Altitude 3 | 0.19 ± 0.15 | | 0.51 ± 0.33 | 0.37 ± 0.44 | 0.18 ± 0.14 | 0.16 ± 0.13 |
| Natural | 0.25 ± 0.18 | | 0.57 ± 0.26 | 0.26 ± 0.27 | 0.21 ± 0.11 | 0.16 ± 0.13 |
| Fragmented | 0.25 ± 0.18 | | 0.61 ± 0.29 | 0.26 ± 0.27 | 0.22 ± 0.15 | 0.16 ± 0.13 |
| TOTAL | 0.25 ± 0.18 | | 0.59 ± 0.27 | 0.22 ± 0.22 | 0.21 ± 0.13 | 0.15 ± 0.11 |

**Table 5:** Means and standard deviations of RMSE for error of point count-based, transect-based, and combined log relative abundance estimators, for individual regions and grouped by altitude, environment type, and total.

| | $\widehat{\ell}_Z$ rel. err. | $\widehat{\ell}_X$ rel. err. | | $\widehat{\ell}_\beta$ rel. err. | |
| --- | --- | --- | --- | --- | --- |
| | (mean ± std) | mean $\bar{p}_b$ | exact $p_b$ | mean $\bar{p}_b$ | exact $p_b$ |
| Bombuscaro | 0.26 ± 0.15 | 0.53 ± 0.15 | 0.18 ± 0.13 | 0.22 ± 0.11 | 0.14 ± 0.09 |
| Copalinga | 0.29 ± 0.17 | 0.58 ± 0.18 | 0.23 ± 0.16 | 0.25 ± 0.18 | 0.13 ± 0.11 |
| ECSF | 0.24 ± 0.12 | 0.63 ± 0.20 | 0.22 ± 0.13 | 0.22 ± 0.10 | 0.16 ± 0.09 |
| Finca | 0.17 ± 0.11 | 0.52 ± 0.18 | 0.16 ± 0.14 | 0.16 ± 0.10 | 0.12 ± 0.09 |
| Cajanuma | 0.20 ± 0.14 | 0.33 ± 0.20 | 0.15 ± 0.10 | 0.17 ± 0.11 | 0.12 ± 0.08 |
| Bellavista | 0.21 ± 0.18 | 0.53 ± 0.11 | 0.34 ± 0.20 | 0.18 ± 0.12 | 0.17 ± 0.12 |
| Altitude 1 | 0.28 ± 0.16 | 0.56 ± 0.17 | 0.21 ± 0.15 | 0.24 ± 0.12 | 0.16 ± 0.11 |
| Altitude 2 | 0.21 ± 0.12 | 0.58 ± 0.20 | 0.19 ± 0.13 | 0.20 ± 0.10 | 0.14 ± 0.09 |
| Altitude 3 | 0.21 ± 0.16 | 0.46 ± 0.17 | 0.17 ± 0.19 | 0.18 ± 0.12 | 0.15 ± 0.11 |
| Natural | 0.25 ± 0.14 | 0.55 ± 0.20 | 0.19 ± 0.13 | 0.22 ± 0.11 | 0.15 ± 0.09 |
| Fragmented | 0.25 ± 0.17 | 0.56 ± 0.17 | 0.23 ± 0.17 | 0.22 ± 0.13 | 0.16 ± 0.11 |
| Total | 0.25 ± 0.15 | 0.57 ± 0.22 | 0.23 ± 0.22 | 0.22 ± 0.12 | 0.15 ± 0.10 |

**Figure 6:**
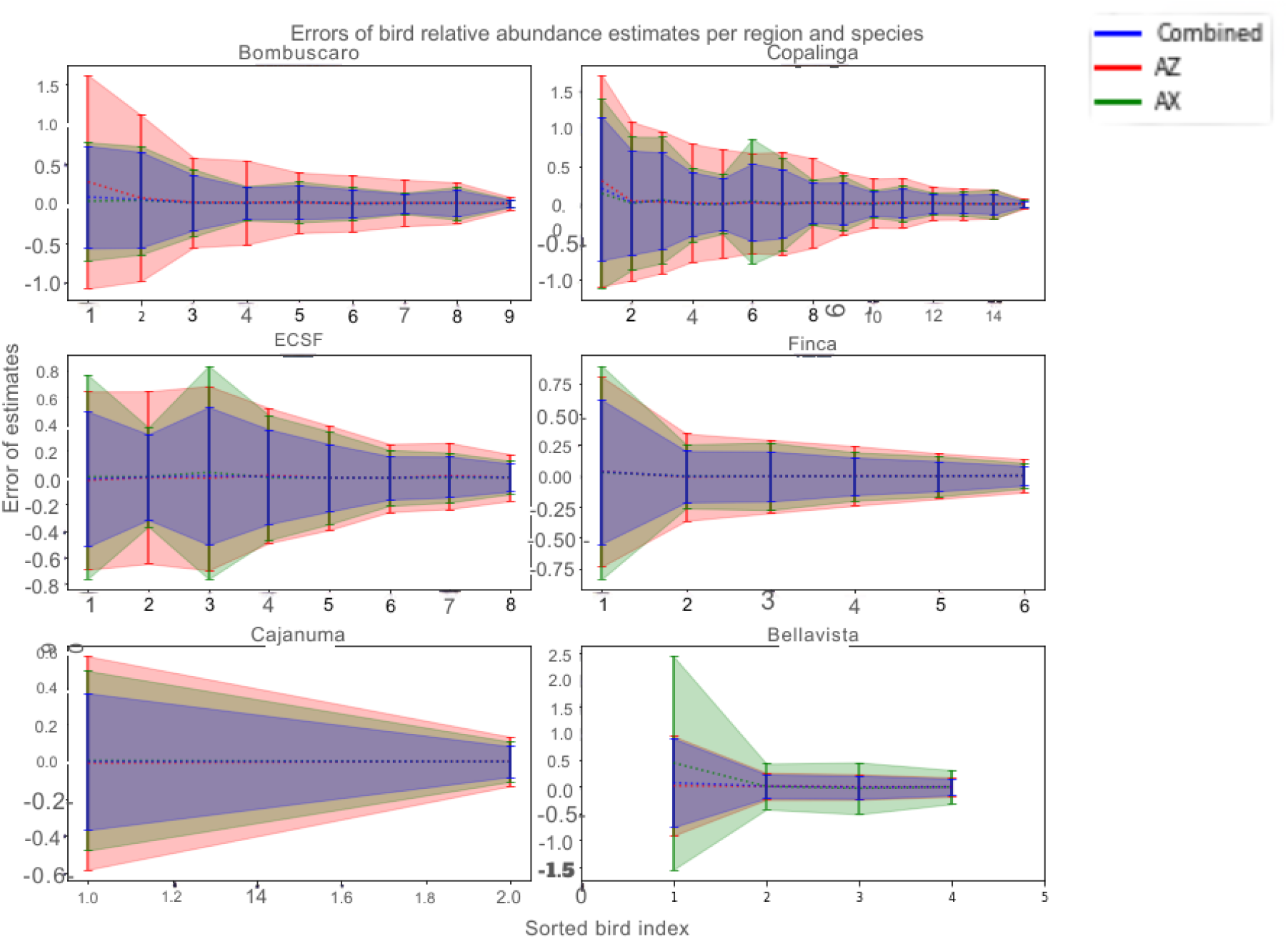
As Figure 5, but with the transect-based and combined estimators using the exact bird weight exponent *p*_*b,n*_ (case b). Confidence intervals for *AX* and the combined estimator are substantially narrower than in Figure 5, particularly for the transect estimator, demonstrating the large benefit of accurate knowledge of the weight-energy relationship.

Figures B8 and C9 of Supplement B and Supplement C respectively show similar results for relative errors of different estimators of log relative abundance. Taken together, the results indicate that more accurate knowledge about individual bird species’ energy consumption habits and requirements can enable improved estimates of relative abundance and log relative abundance. The degree of improvement is displayed quantitatively in Table 4 (for relative abundance estimators) and in Table 5 (for log relative abundance estimators).

## 5 Discussion

### 5.1 Fruit weight and bird weight as predictors of energy consumption

A central finding of this study is that mean fruit weight consumed does not significantly influence the number of feeding observations, once bird weight is accounted for. This result held consistently across 11 of the 12 data subsets examined (individual regions, elevational groups, habitat types, and the combined dataset), and was corroborated by the systematic reduction in AIC values when the fruit weight parameter *p*_*f*_ was dropped from the model. This finding contrasts with the intuition that larger fruit should require fewer foraging bouts due to higher energy yield per item, but is consistent with field observations suggesting that frugivores often adjust bite size rather than foraging frequency when consuming fruit of varying sizes [7]. It also aligns with the well-established view that frugivore foraging rates are constrained primarily by digestive processing time, which scales with gut volume and hence with bird body size rather than with fruit size per se [8].

The bird weight exponent *p*_*b*_, by contrast, was estimated to be consistently negative (overall 95% confidence interval [−0.6, − 0.1]), indicating that heavier bird families have fewer transect feeding observations per individual relative to what their point count abundance would predict. As discussed in Section 4, this is consistent with *p*_*b*1_ ≈ 1, implying that energy consumption per feeding event scales approximately linearly with bird body mass. This interpretation aligns with prior allometric studies reporting that basal metabolic rate scales with body mass raised to a power near 0.7 for most avian families [14], while absolute food intake per bout scales more steeply. The larger confidence intervals observed at low elevations, relative to higher elevations, are consistent with the greater phylogenetic and body-size diversity of frugivore assemblages in warmer lowland forests, which would introduce more variability in the relationship between body size and foraging rate. Future work using species-level body mass data, rather than family-level averages, could reduce this variability and yield tighter parameter estimates.

### 5.2 Combining point counts and transect observations for improved abundance estimation

The simulation results demonstrate that point count-based and transect observation-based estimators of relative abundance are complementary in their error structure, with the point count estimator consistently showing lower variance under the conditions simulated. The negative correlation between the feeding-to-observation ratio and the number of transect observations (Figure 3) provides empirical confirmation of regression to the mean in the transect-based estimates, which is the mechanism that makes the two estimator types counterbalancing. This counterbalancing effect is precisely what the linear combination exploits: by assigning weight *β* to the point count estimator and 1 − *β* to the transect estimator, the combined estimator achieves a variance lower than either component alone, as guaranteed by the minimum-variance formula (30).

The practical implication is that field studies collecting both point count and transect data need not choose between them at the analysis stage. Instead, both sources can be integrated using the framework developed here to obtain abundance estimates with formally smaller uncertainty. The improvement is most pronounced when the investigator has accurate prior knowledge of the bird weight exponent *p*_*b,n*_: in this case, the transect-based estimator becomes substantially more informative, and the combined estimator achieves relative RMSE values roughly 40% lower than the point count estimator alone (Table 4). When only the mean value 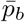 is available, the improvement is more modest but still consistent across all regions and groupings. This suggests that investing effort in characterizing the allometric energy scaling of the focal bird families— whether from prior literature or from empirical calibration within the study system—pays dividends in the accuracy of final abundance estimates.

These findings have direct relevance for ongoing monitoring programs in Andean forests, where both survey methods are commonly deployed but rarely integrated analytically. The Podocarpus region in particular supports well-documented gradients in frugivore community composition across elevation and habitat fragmentation [6, 16], and the present framework could be applied to quantify how relative abundance of key seed-dispersing families changes across these gradients with greater precision than point counts alone would allow. Similar applications are conceivable in any system where ecologists collect concurrent point count and behavioral observation data, including insectivore foraging studies and waterbird monitoring programs.

### 5.3 Confidence intervals and their calibration

A notable practical contribution of this paper is the bootstrapping procedure for computing confidence intervals for the combined abundance estimator. Table 6 shows that the 90% confidence intervals produced by this procedure contain the true relative abundance in approximately 90% of simulation instances across all regions, groupings, and parameterizations tested. This calibration is remarkably consistent, with proportions ranging from 0.900 to 0.908 regardless of whether the mean or exact bird weight exponent was used. The robustness of the calibration across these two scenarios suggests that the bootstrapping procedure is not highly sensitive to uncertainty in the weight exponent, which is an important practical property since this parameter must typically be estimated from data.

**Table 6:**
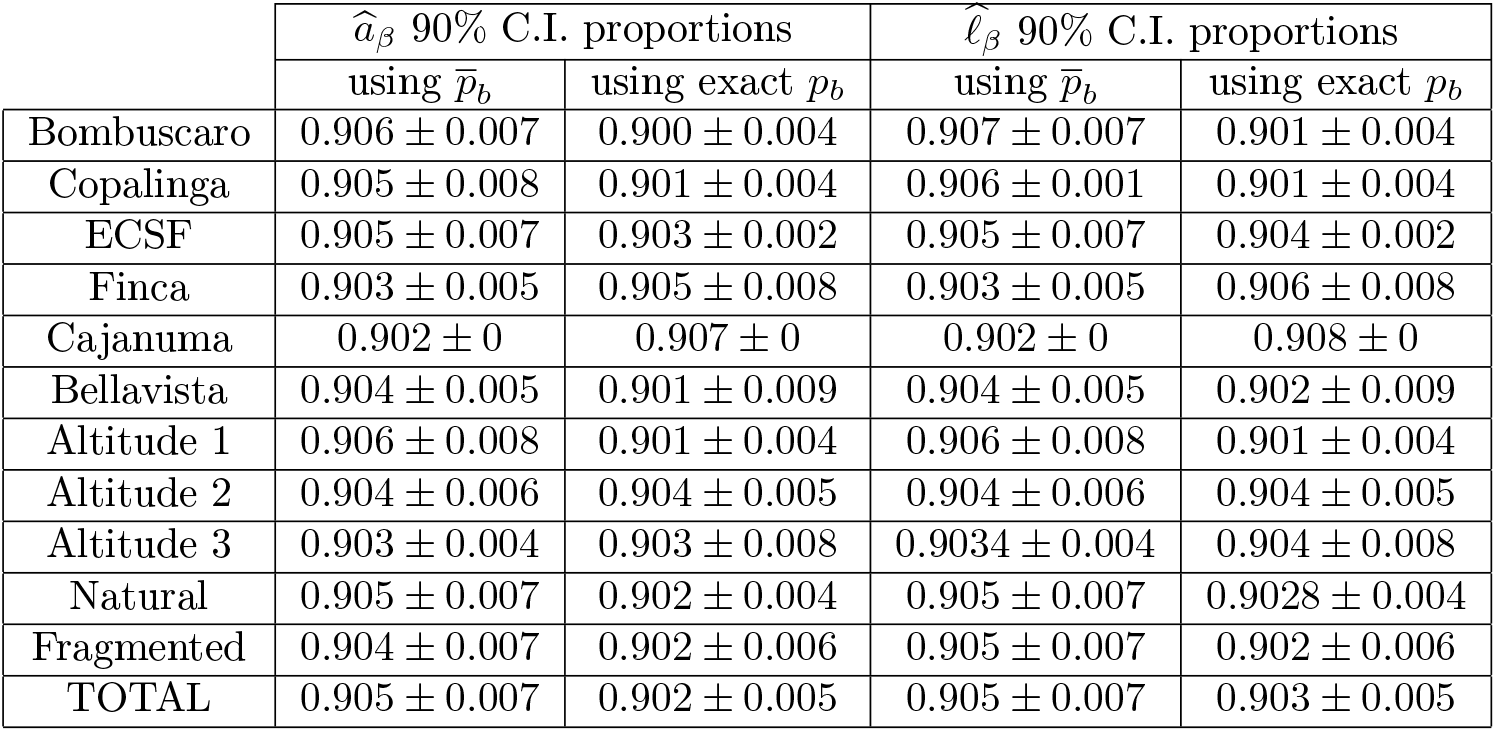
Proportion of instances where the true relative abundance (respectively log relative abundance) lay within the 90% confidence intervals computed by the bootstrapping technique described in Section 3.1.7. Estimates using the mean value of the bird weight exponent 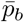 and the exact value of *p*_*b*_ are shown. The error bars are for one standard deviation calculated from the proportions obtained for different families in the region. Proportions consistently near 0.90 across all regions confirm that the bootstrapping procedure produces well-calibrated confidence intervals.

| | $\widehat{a}_\beta$ 90% C.I. proportions | | $\widehat{\ell}_\beta$ 90% C.I. proportions | |
| --- | --- | --- | --- | --- |
| | using $\bar{p}_b$ | using exact $p_b$ | using $\bar{p}_b$ | using exact $p_b$ |
| Bombuscaro | 0.906 ± 0.007 | 0.900 ± 0.004 | 0.907 ± 0.007 | 0.901 ± 0.004 |
| Copalinga | 0.905 ± 0.008 | 0.901 ± 0.004 | 0.906 ± 0.001 | 0.901 ± 0.004 |
| ECSF | 0.905 ± 0.007 | 0.903 ± 0.002 | 0.905 ± 0.007 | 0.904 ± 0.002 |
| Finca | 0.903 ± 0.005 | 0.905 ± 0.008 | 0.903 ± 0.005 | 0.906 ± 0.008 |
| Cajanuma | 0.902 ± 0 | 0.907 ± 0 | 0.902 ± 0 | 0.908 ± 0 |
| Bellavista | 0.904 ± 0.005 | 0.901 ± 0.009 | 0.904 ± 0.005 | 0.902 ± 0.009 |
| Altitude 1 | 0.906 ± 0.008 | 0.901 ± 0.004 | 0.906 ± 0.008 | 0.901 ± 0.004 |
| Altitude 2 | 0.904 ± 0.006 | 0.904 ± 0.005 | 0.904 ± 0.006 | 0.904 ± 0.005 |
| Altitude 3 | 0.903 ± 0.004 | 0.903 ± 0.008 | 0.9034 ± 0.004 | 0.904 ± 0.008 |
| Natural | 0.905 ± 0.007 | 0.902 ± 0.004 | 0.905 ± 0.007 | 0.9028 ± 0.004 |
| Fragmented | 0.904 ± 0.007 | 0.902 ± 0.006 | 0.905 ± 0.007 | 0.902 ± 0.006 |
| TOTAL | 0.905 ± 0.007 | 0.902 ± 0.005 | 0.905 ± 0.007 | 0.903 ± 0.005 |

Rigorous confidence intervals for abundance estimates are rarely reported in avian community monitoring studies, which tend to rely on informal descriptions of survey effort as a proxy for precision. The present framework offers a path toward more formal uncertainty quantification that is grounded in the statistical properties of the underlying observation process. This is particularly valuable in the context of long-term monitoring, where changes in abundance over time must be distinguished from sampling variability—a distinction that requires well-calibrated confidence intervals.

### 5.4 Limitations and future directions

Several limitations of the present study should be acknowledged. First, the theoretical framework assumes that point count and transect observations are made independently and that the detection probability within the specified radius or distance band is constant across species and habitats. Violations of the constant detection probability assumption—which are common in heterogeneous landscapes—could bias both the point count and transect estimators in ways not captured by the current model. Distance sampling methods [10] or *N* -mixture models could in principle be integrated into the framework to relax this assumption.

Second, the empirical application combines observations across bird families, which span a wide range of body sizes and behavioral repertoires. Family-level aggregation, while necessary given the relatively small per-species sample sizes in the dataset, smooths over ecologically meaningful interspecific variation. Future applications with larger datasets could operate at the species or genus level, potentially yielding more informative estimates of *p*_*b,n*_ and hence more accurate transect-based abundance estimators.

Third, the simulation-based evaluation uses the observed data as “ground truth,” which means the simulations inherit whatever systematic biases may be present in the original field data. Independent validation using replicated field datasets from different sites or time periods would provide a stronger test of the framework’s generalizability.

Despite these limitations, the methods developed here represent a step toward more rigorous integration of complementary survey data types in avian ecology, and the general approach of linking energy consumption models to multiple observation processes could be extended to other taxa and monitoring contexts.

## 6 Conclusions

The principal conclusions of this study are as follows:

- For the dataset drawn from six habitats in Ecuador described in Section 3.2.1, there is no statistical evidence that bird species’ energy consumption depends on the weight of fruit consumed.
- On the other hand, there is statistical evidence from this dataset supporting existing literature that energy consumption of bird species depends on bird weight.
- There is also statistical evidence from this same dataset that energy consumption per transect observation is larger for heavier birds than for lighter birds.
- Simulations based on the Ecuador data demonstrate that point counts and transect observations can be used to obtain independent estimates of relative abundance and log of relative abundance. Typically, the point count-based estimates are more accurate.
- The two estimates mentioned in the previous item may be linearly combined to form another estimate which has lower RMSE. The user may run simulations to estimate the parameter *β* needed to form a linear combination of point count based and transect observation based abundance estimators with smaller root mean squared error.
- Error bars may be computed for the linearly combined relative abundance estimates that accurately reflect the uncertainty in these estimates.
- With more accurate information concerning the dependence of bird type energy consumption as a function of weight, a better estimate of *β* can be made that leads to combined estimators of abundance with lower root mean squared error.

## CRediT statement

**Conceptualization**: V.S., M.J.A.B., A.B., C.T.; **Data curation**: V.S., M.Q., E.L.N., M.S.; **Formal analysis**: M.J.A.B., A.B., C.T.; **Funding acquisition**: K.B.S., O.A.; **Investigation**: V.S., M.Q., E.L.N., M.S.; **Methodology**: M.J.A.B., A.B., C.T.; **Project administration**: V.S., K.B.S., O.A.; **Resources**: K.B.S., O.A.; **Software**: M.J.A.B., A.B., C.T.; **Supervision**: K.B.S., V.S., C.T.; **Validation**: M.J.A.B., C.T.; **Visualization**: M.J.A.B., A.B., C.T.; **Writing – original draft**: M.J.A.B., A.B., C.T.; **Writing – review & editing**: V.S., E.L.N., C.T.

## Acknowledgements

We are grateful to the Fulbright Program administered by the US Department of State, the Catholic University of Cuenca and to Texas A&M University-Central Texas for providing travel funding and logistical support during the conduct of this research.

## Supplement A: Model fit diagnostics

This supplement contains scatter plots of actual versus predicted transect observations for the two-coefficient model (Figures A1–A4) and the one-coefficient model (Figures A5–A8), organized by location, elevation, habitat type, and combined.

**Figure A1:**
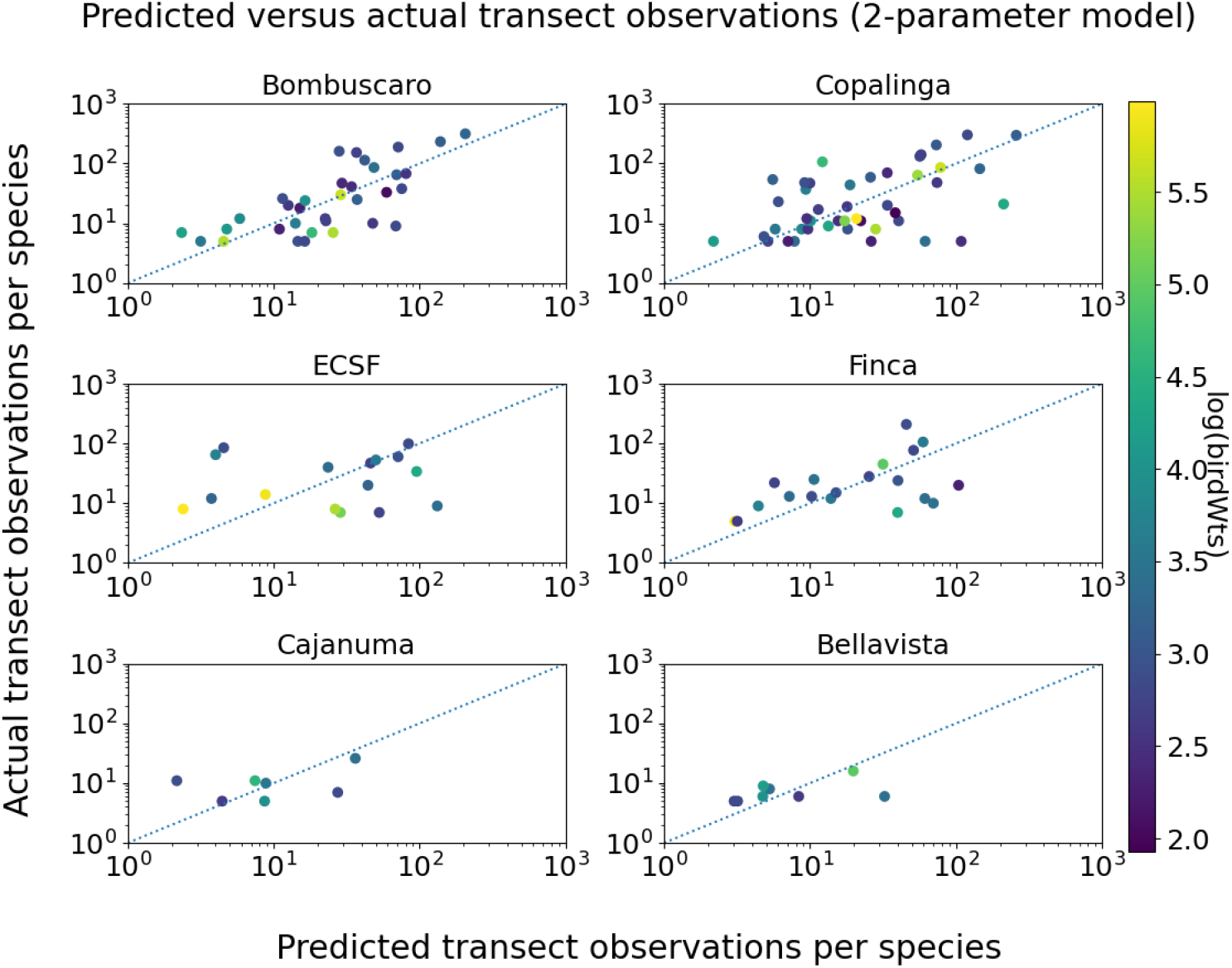
Transect observation prediction accuracy by location for the two-coefficient model (13) that includes both bird weight and fruit weight as predictors. Each panel shows log-scale actual versus log-scale predicted transect observations for one of the six study sites, with point color indicating mean bird species weight. The 45-degree line corresponds to zero prediction error. Scatter around the line reflects residual variability not explained by bird weight or fruit weight.

**Figure A2:**
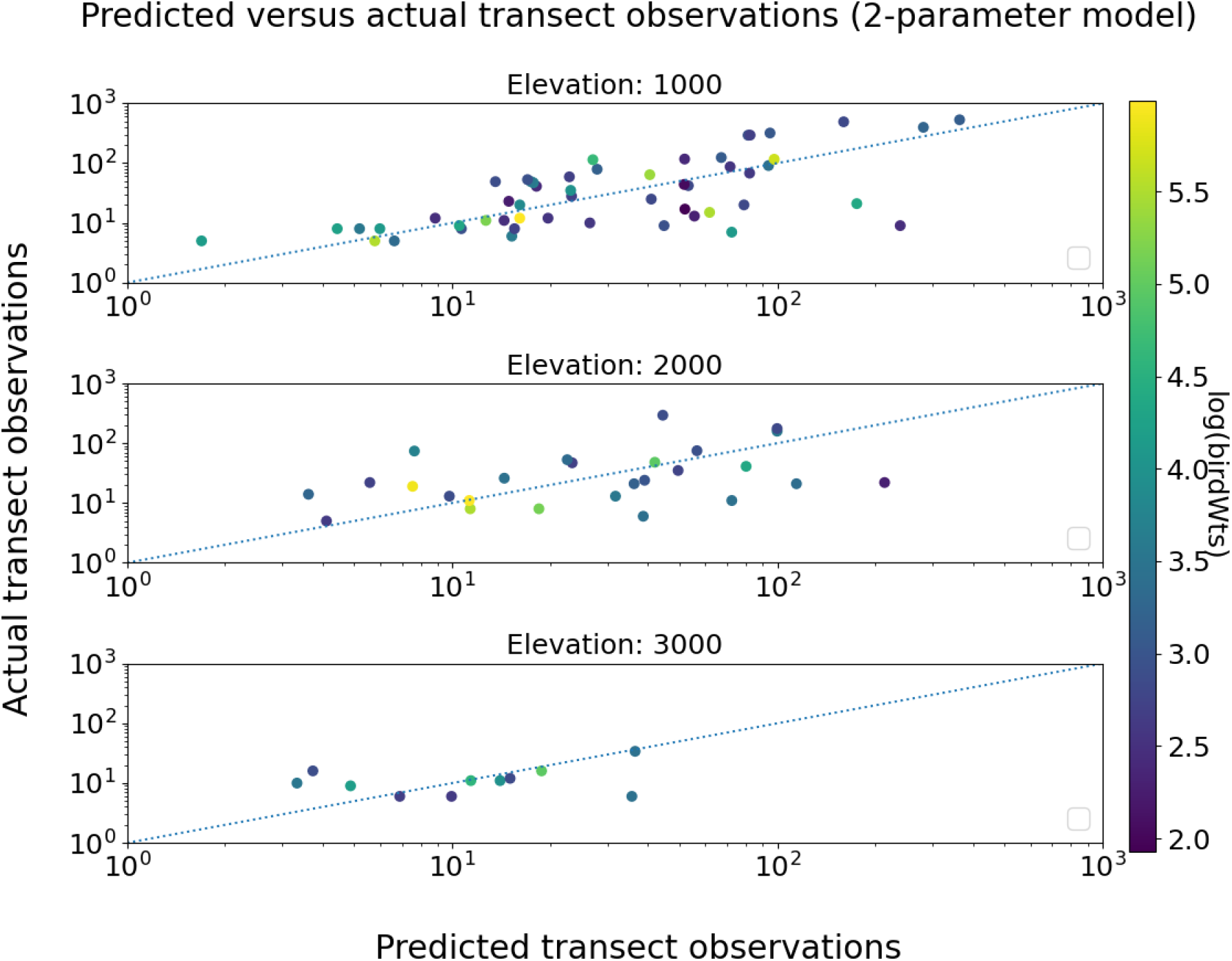
Transect observation prediction accuracy of the two-coefficient model (13) (bird weight + fruit weight), with species data pooled by elevation. Low (1000 m), mid (2000 m), and high (3000 m) elevation panels each combine data from the two sites at that elevation. Axes and line as in Figure A1.

**Figure A3:**
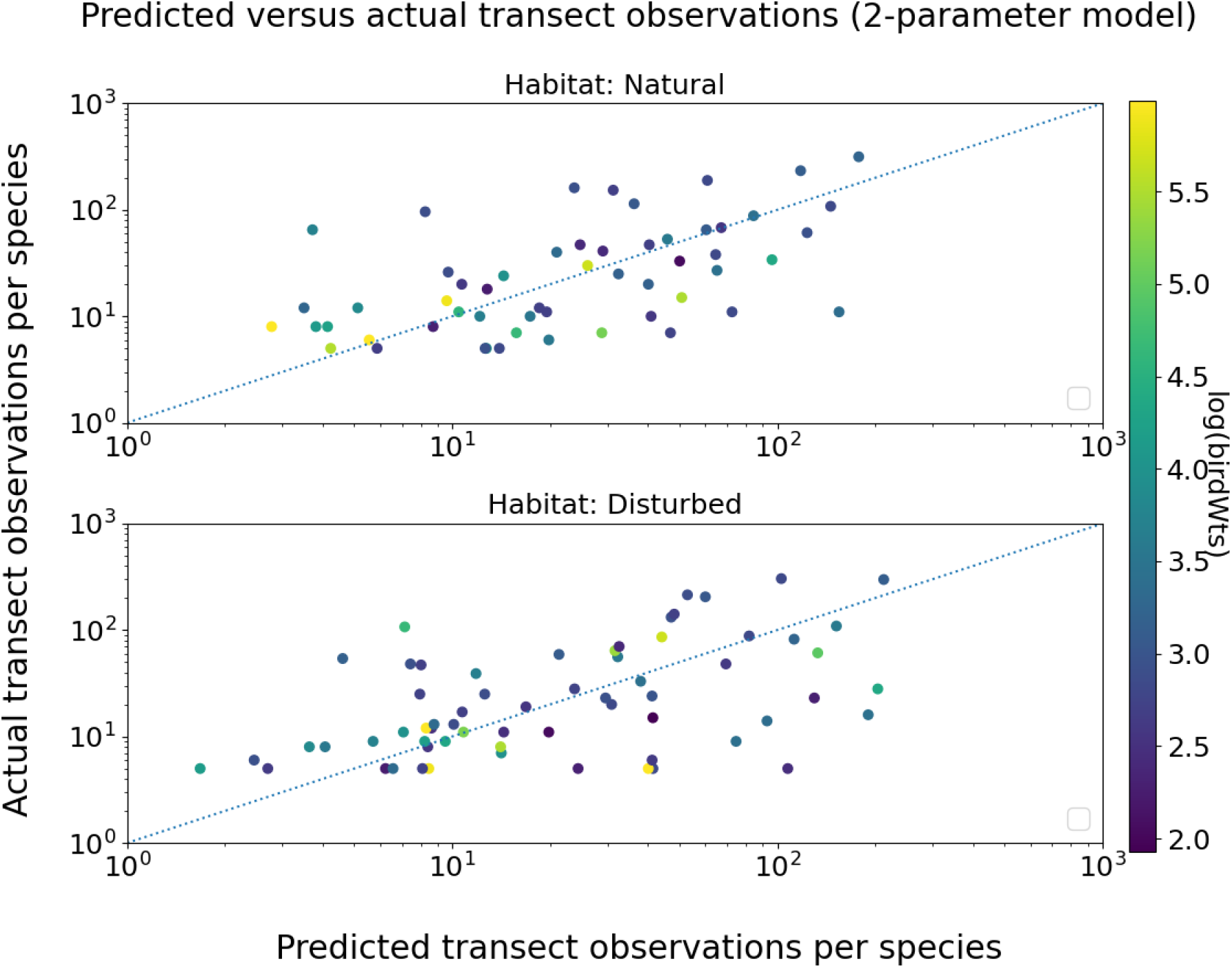
Transect observation prediction accuracy of the two-coefficient model (13) (bird weight + fruit weight), with species data pooled by habitat type. Natural forest and fragmented forest panels each combine data from the three sites of that habitat type. Axes and line as in Figure A1.

**Figure A4:**
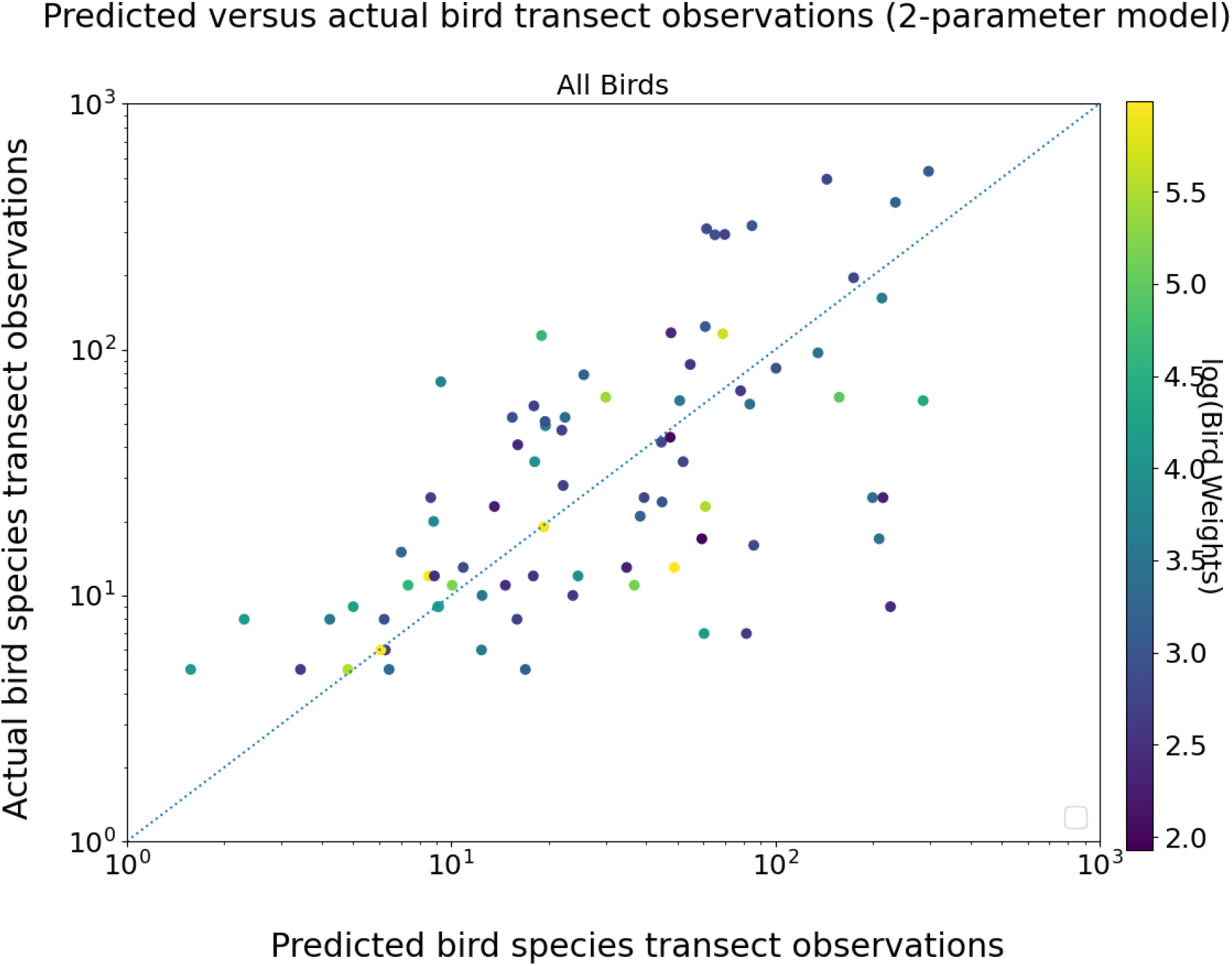
Transect observation prediction accuracy for the two-coefficient model (13) (bird weight + fruit weight) using all species observations from all six sites combined. Axes and line as in Figure A1.

**Figure A5:**
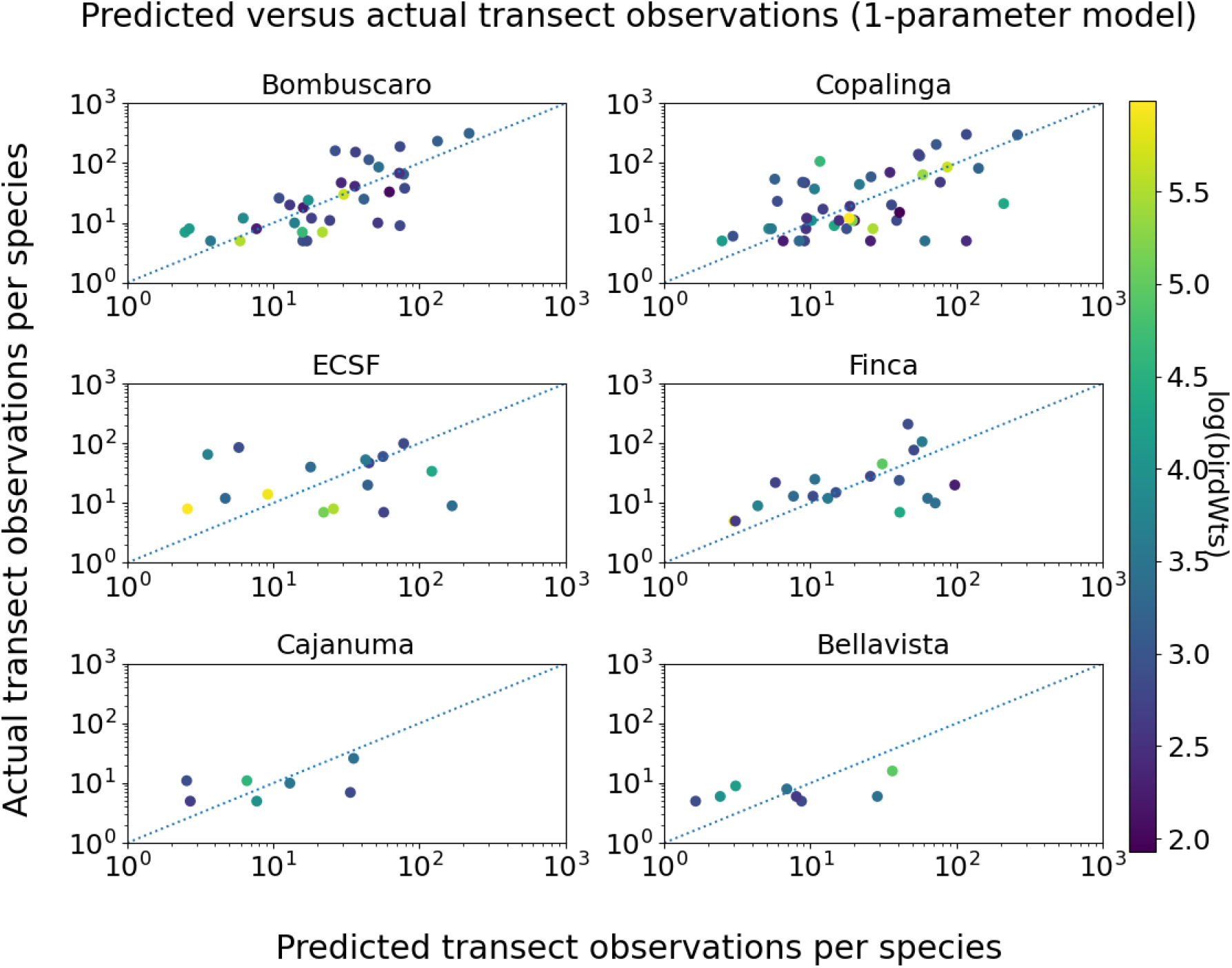
Transect observation prediction accuracy by location for the one-coefficient model (47), which includes bird weight only (*p*_*f*_ = 0). Compare with Figure A1 for the two-coefficient model. Axes and line as in Figure A1.

**Figure A6:**
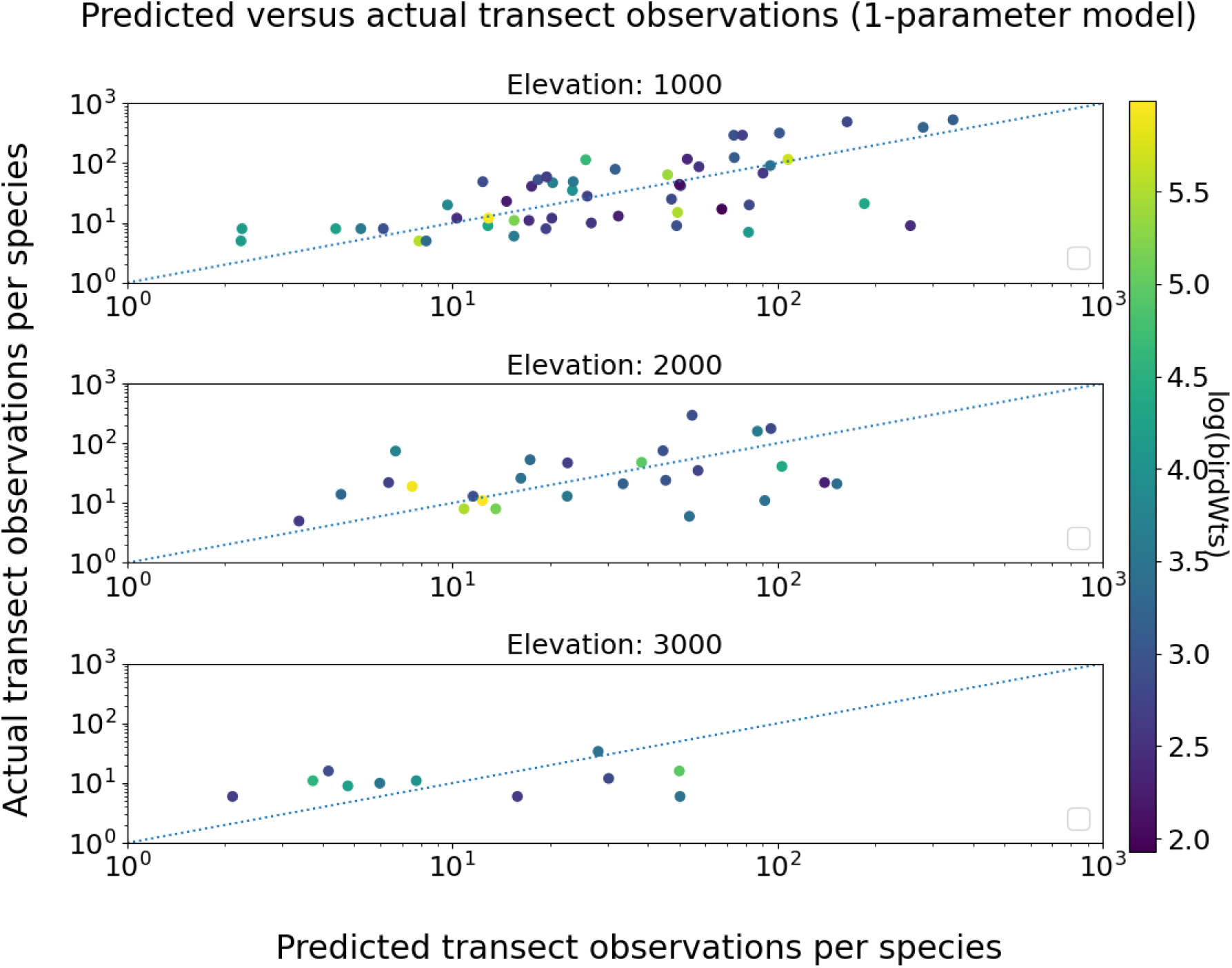
Transect observation prediction accuracy for the one-coefficient model (47) (bird weight only), with species data pooled by elevation. Compare with Figure A2. Axes and line as in Figure A1.

**Figure A7:**
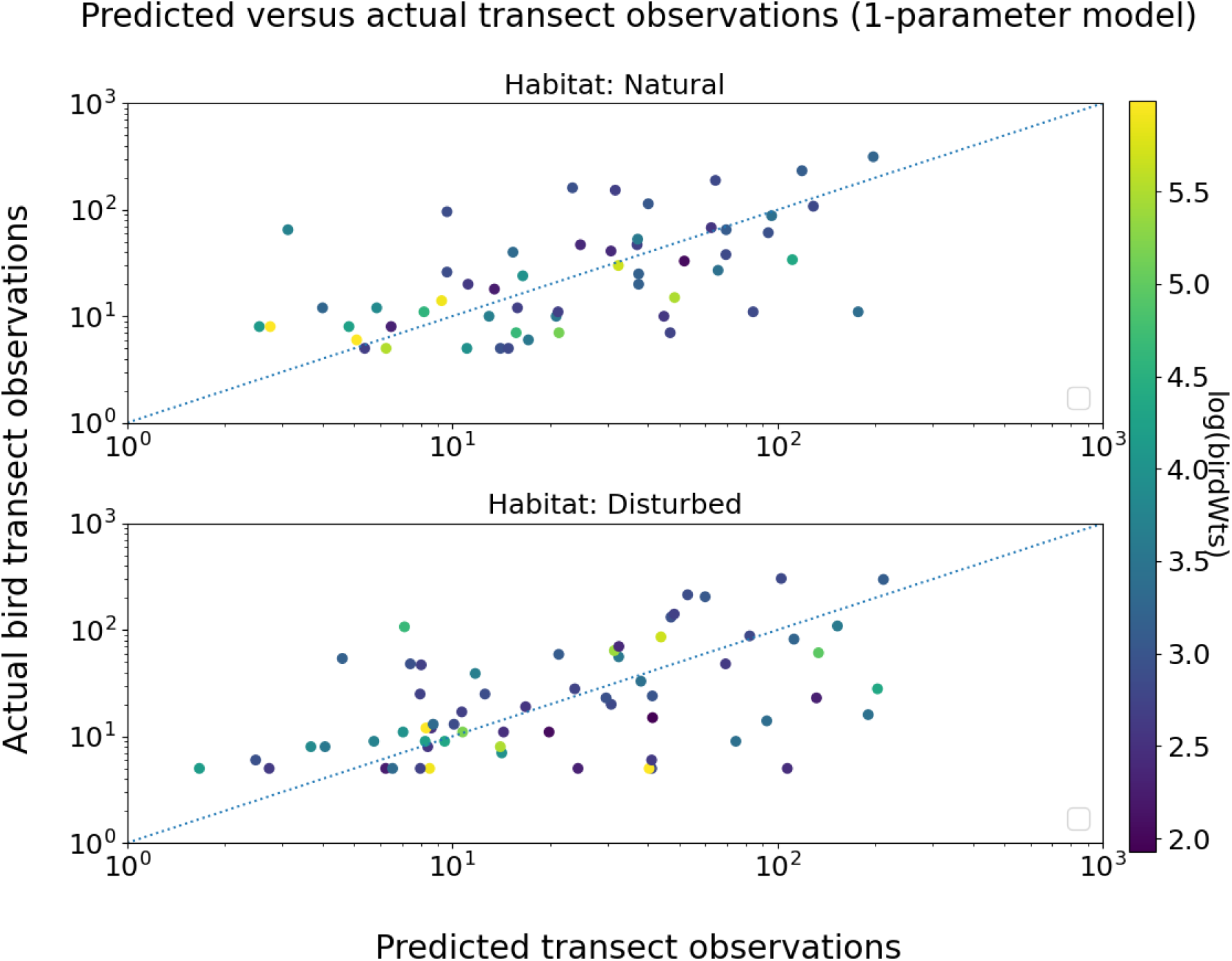
Transect observation prediction accuracy for the one-coefficient model (47) (bird weight only), with species data pooled by habitat type. Compare with Figure A3. Axes and line as in Figure A1.

**Figure A8:**
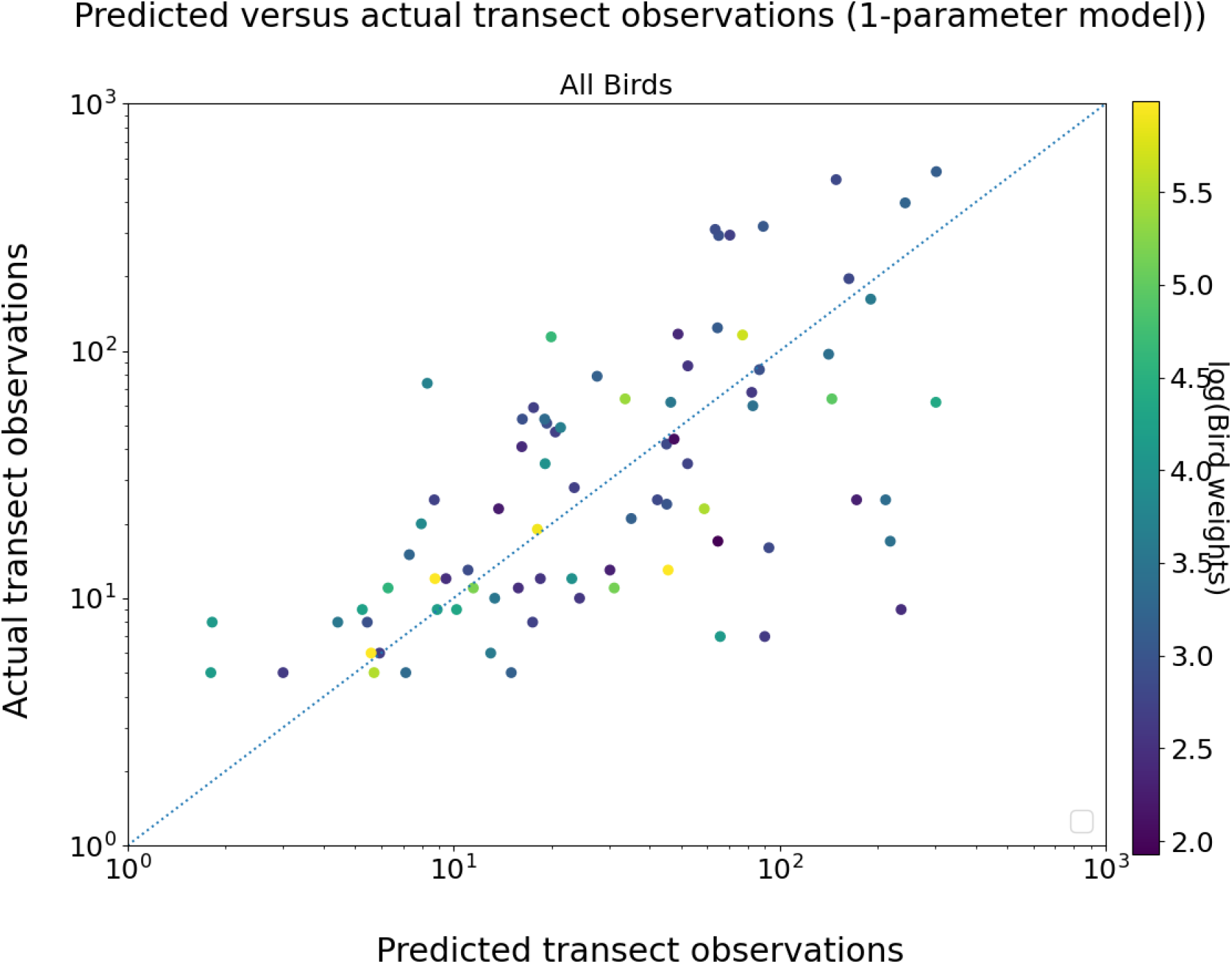
Transect observation prediction accuracy for the one-coefficient model (47) (bird weight only), using all species observations from all six sites combined. Compare with Figure A4. Axes and line as in Figure A1.

## Supplement B: Estimator bias, variance, and RMSE using mean bird weight exponent 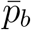 (case a)

This supplement contains detailed performance evaluation figures for the point count-based, transect-based, and combined estimators of relative abundance and log relative abundance, computed using the mean value 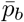 of the bird weight exponent (case a as defined in Section 3.2.3).

**Figure B1:**
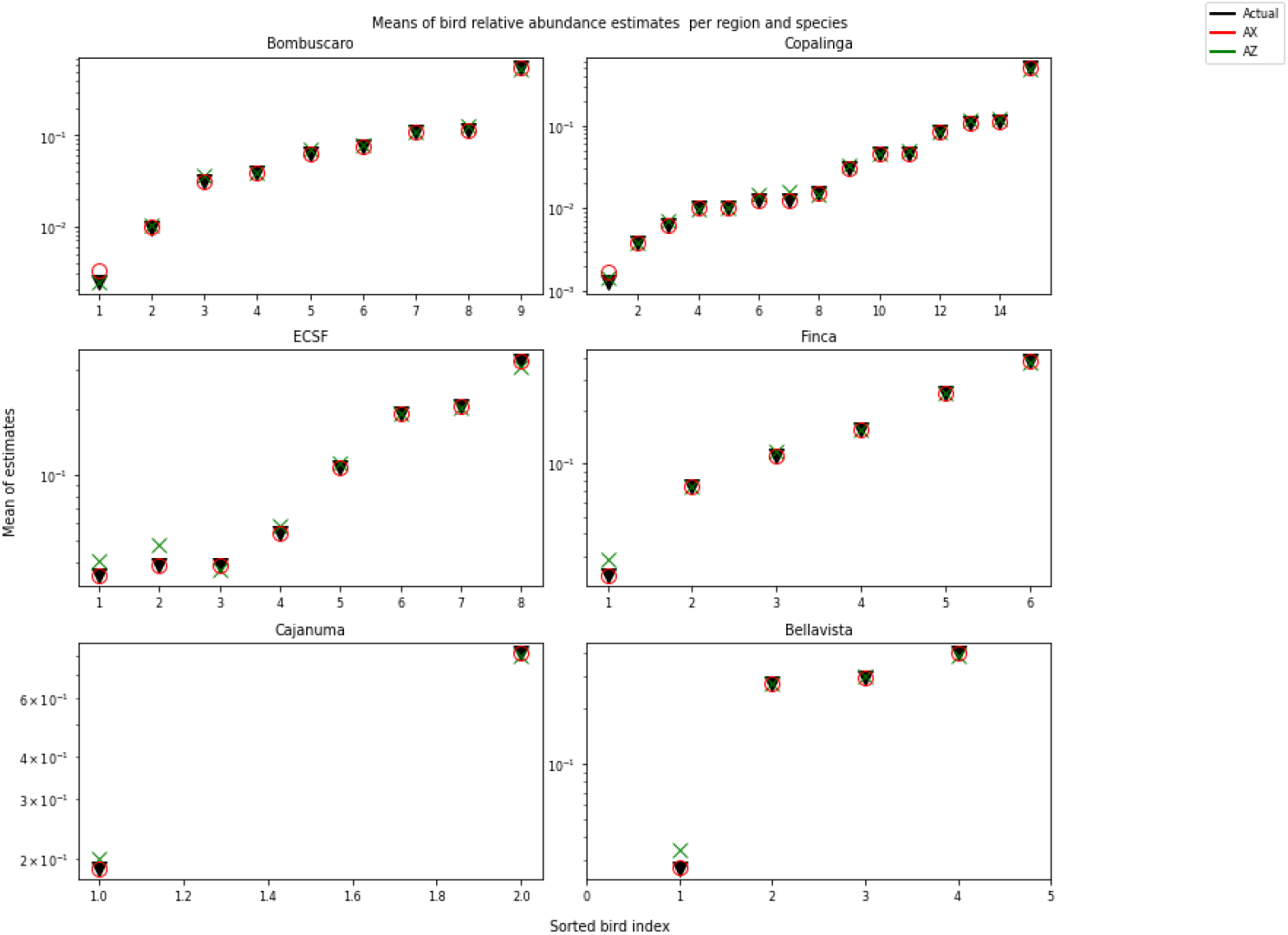
Mean values of the point count-based estimator 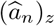 (circles) and the transect-based estimator 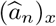 (triangles) over 10,000 simulations, compared to true relative abundance (dashed line), for bird families at each of the six study sites. The transect estimator is computed using the mean bird weight exponent 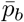 (case a). Within each site panel, families are sorted in order of increasing true relative abundance. Both estimators show little systematic bias, with slightly larger departures for rare families.

**Figure B2:**
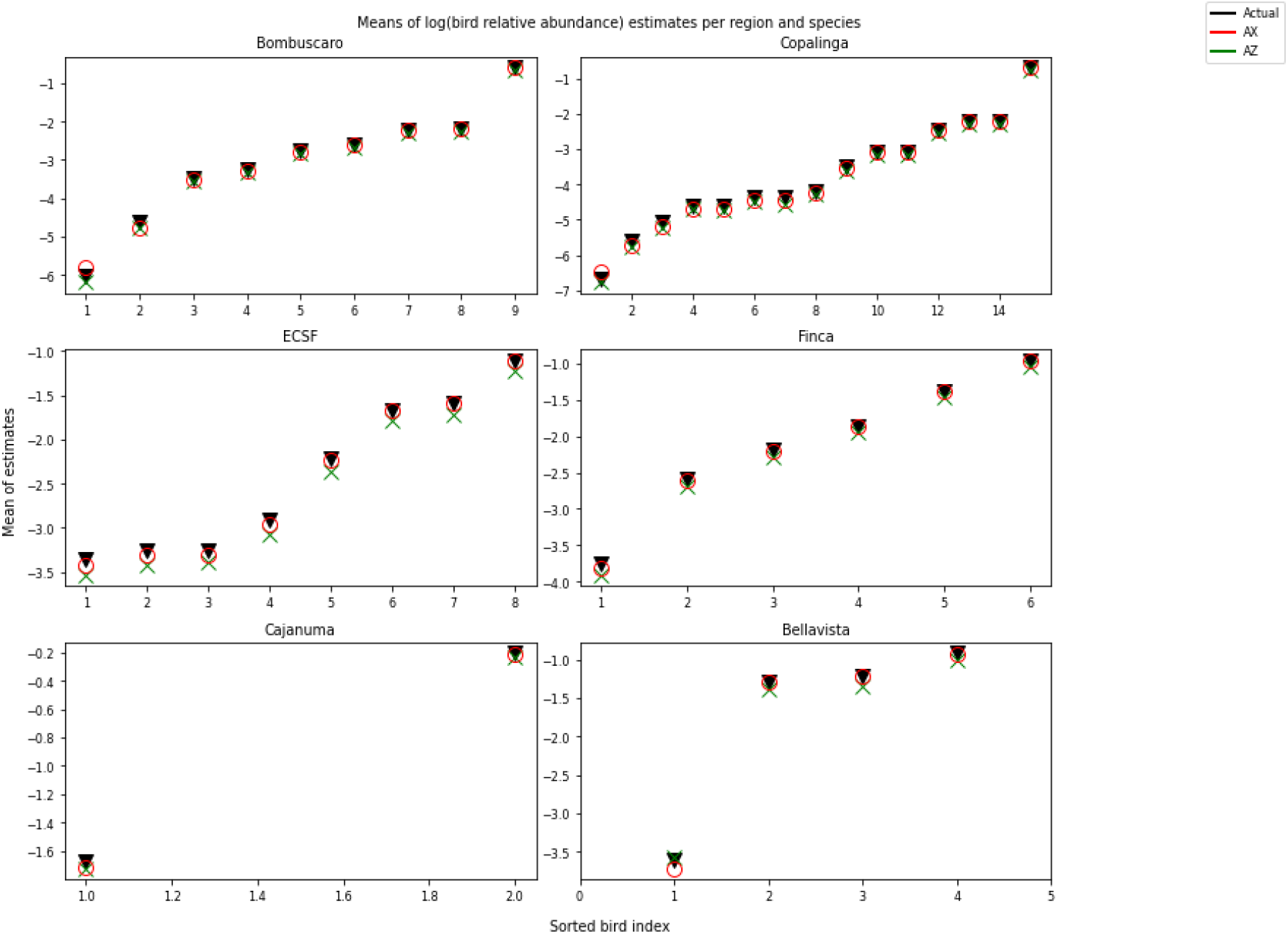
Mean values of the point count-based log abundance estimator 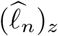 (circles) and the transect-based log abundance estimator 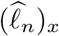 (triangles) over 10,000 simulations, compared to true log relative abundance (dashed line), for bird families at each of the six study sites. The transect estimator uses the mean bird weight exponent 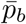 (case a). Families are sorted as in Figure B1.

**Figure B3:**
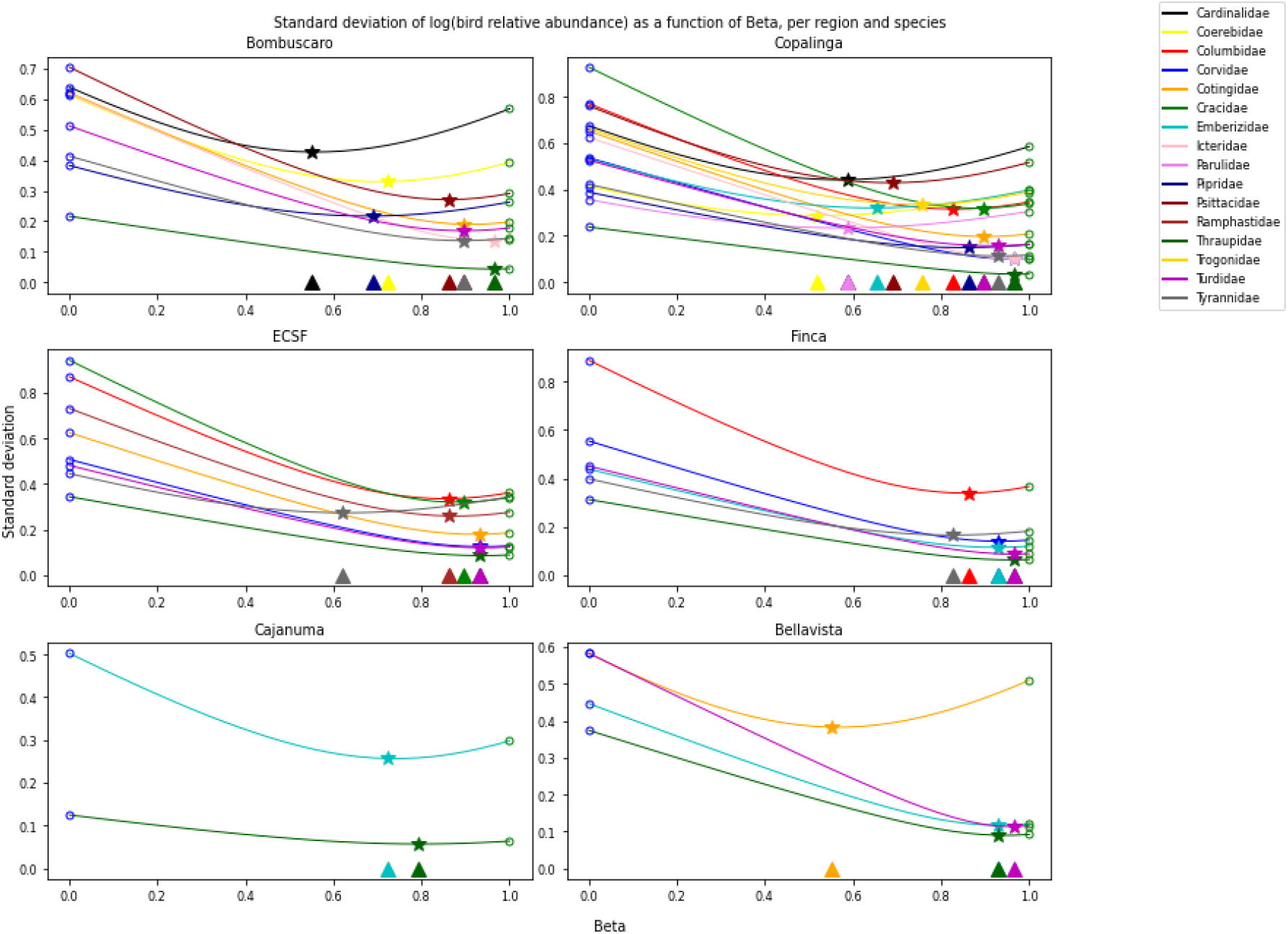
Standard deviation of the combined log abundance estimator 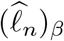 as a function of *β* ∈ [0, 1], for each bird family in each of the six study sites. *β* = 0 corresponds to 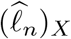 (transect-based); *β* = 1 corresponds to 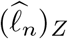 (point count-based). Stars mark the minimum of each curve; triangles on the *x*-axis indicate the theoretically optimal *β* from (35). Transect estimator uses 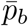 (case a).

**Figure B4:**
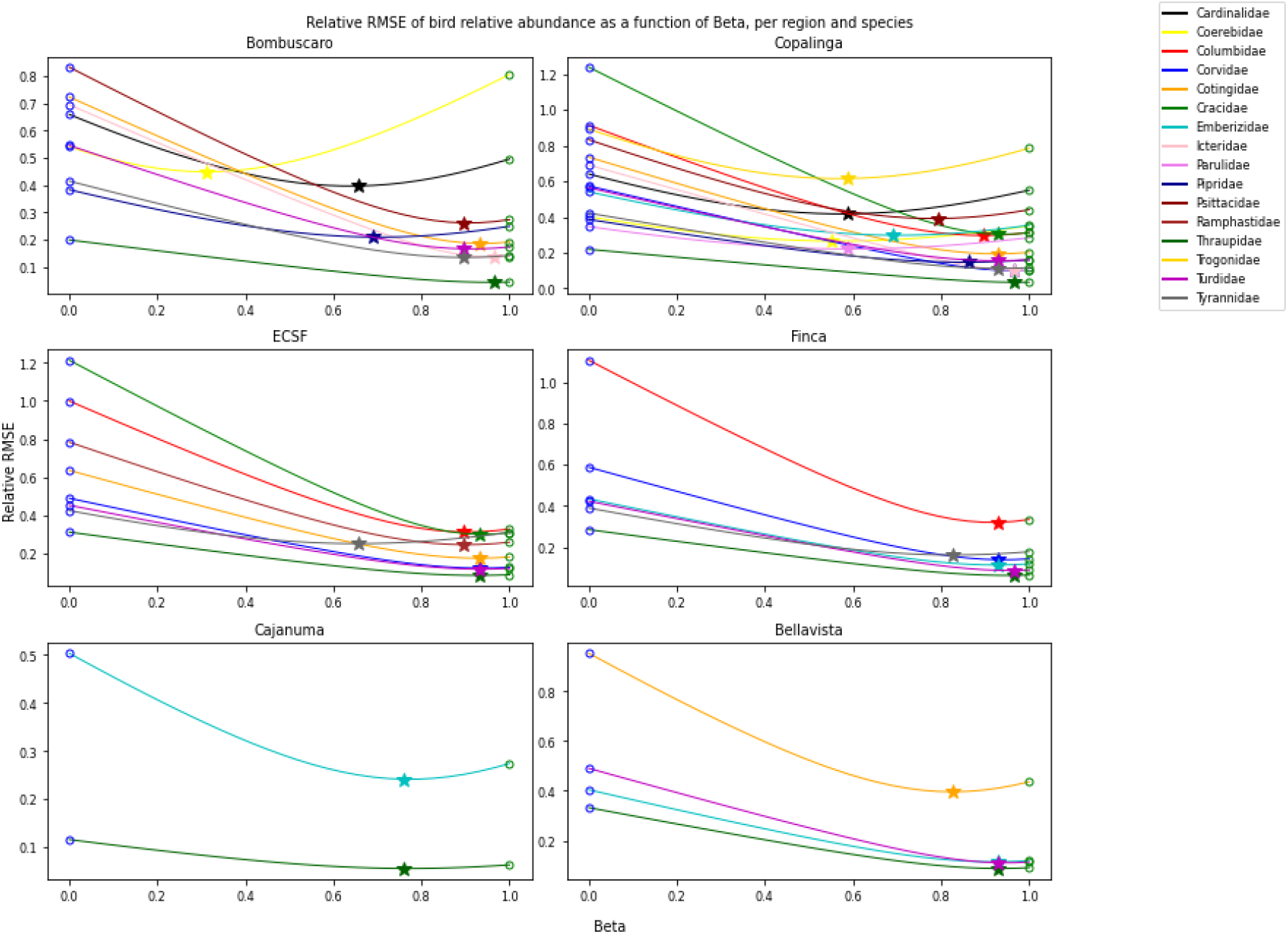
Relative root mean squared error (RMSE divided by true relative abundance) of the combined estimator 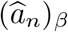 as a function of *β* ∈ [0, 1], for each bird family in each site. *β* = 0 corresponds to 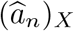; *β* = 1 to 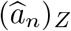. Stars and triangles as in Figure 4 of the main body. The near-identical appearance of this figure and Figure 4 confirms that bias is negligible relative to variance. Transect estimator uses 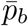 (case a).

**Figure B5:**
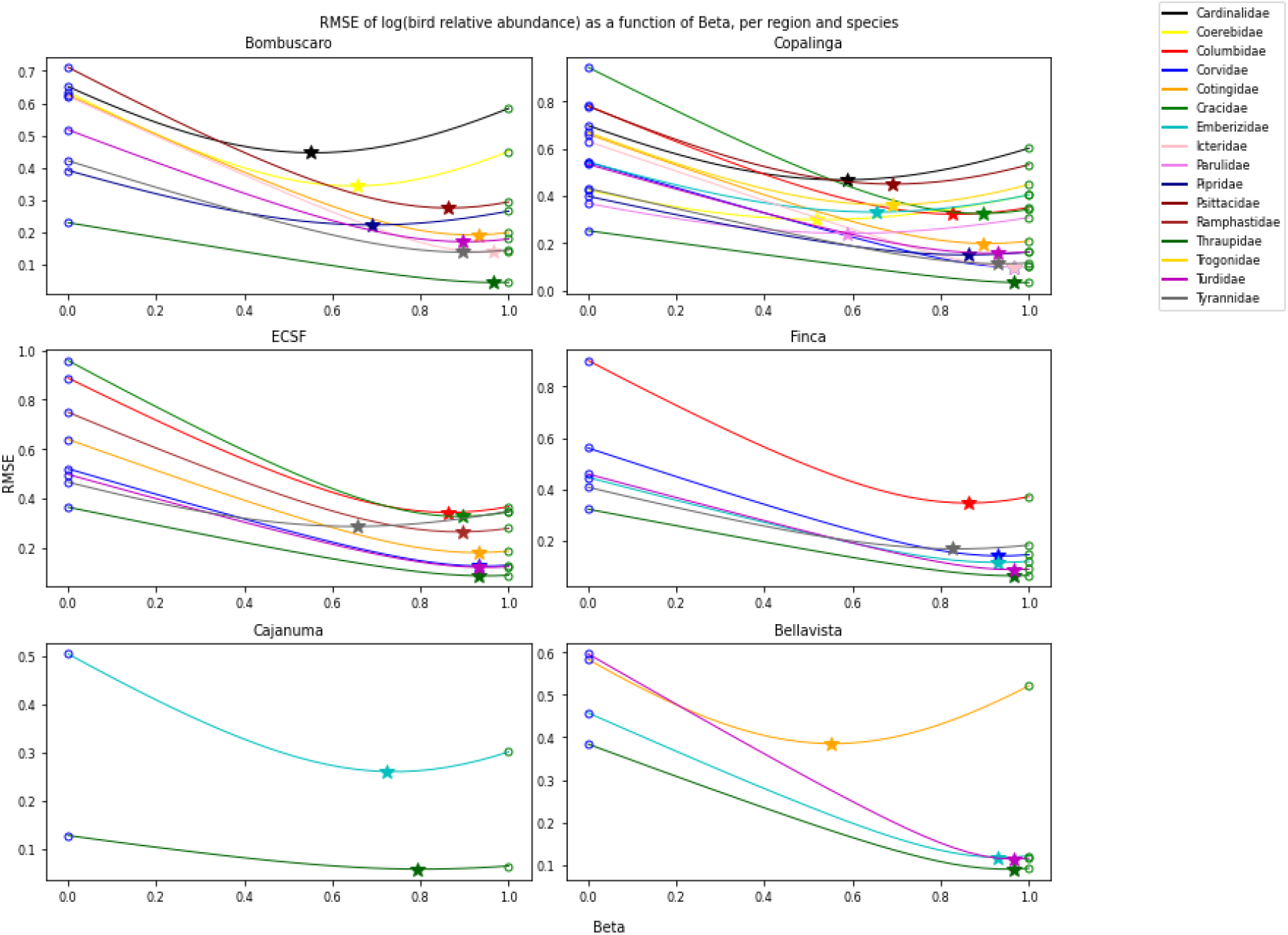
Root mean squared error (RMSE) of the combined log abundance estimator 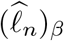 as a function of *β* ∈ [0, 1], for each bird family in each site. *β* = 0 corresponds to 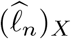; *β* = 1 to 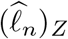. Stars and triangles as in Figure B3. Transect estimator uses 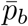 (case a).

**Figure B6:**
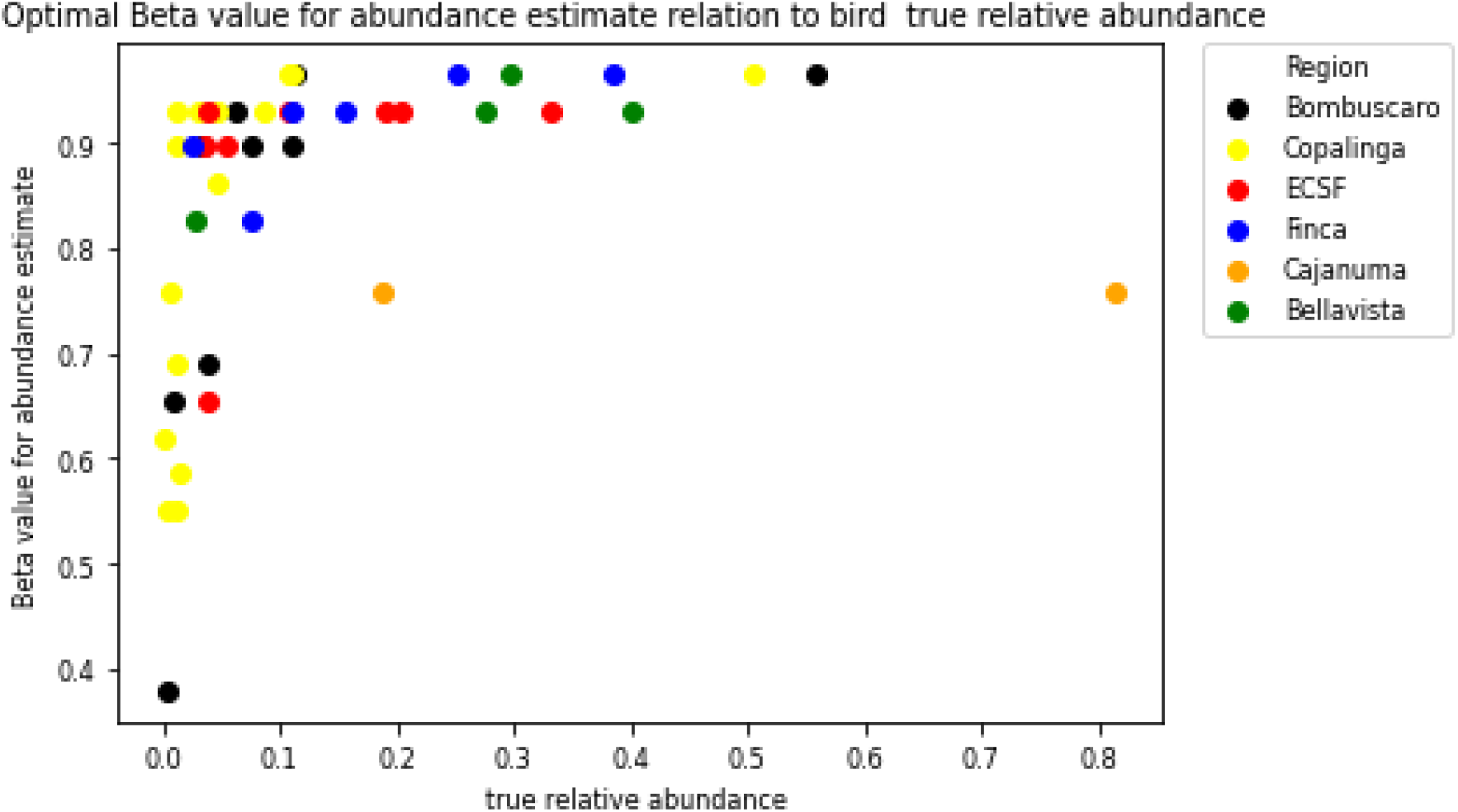
RMSE-minimizing value of *β* for the combined relative abundance estimator 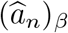, plotted against the true relative abundance of each bird family, for all families across all six sites. Each point represents one family-site combination. Lower *β* values (giving more weight to the transect estimator) occur predominantly for rare families (true relative abundance below 0.1), while common families consistently receive high *β* (closer to the point count estimator). Transect estimator uses 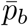 (case a).

**Figure B7:**
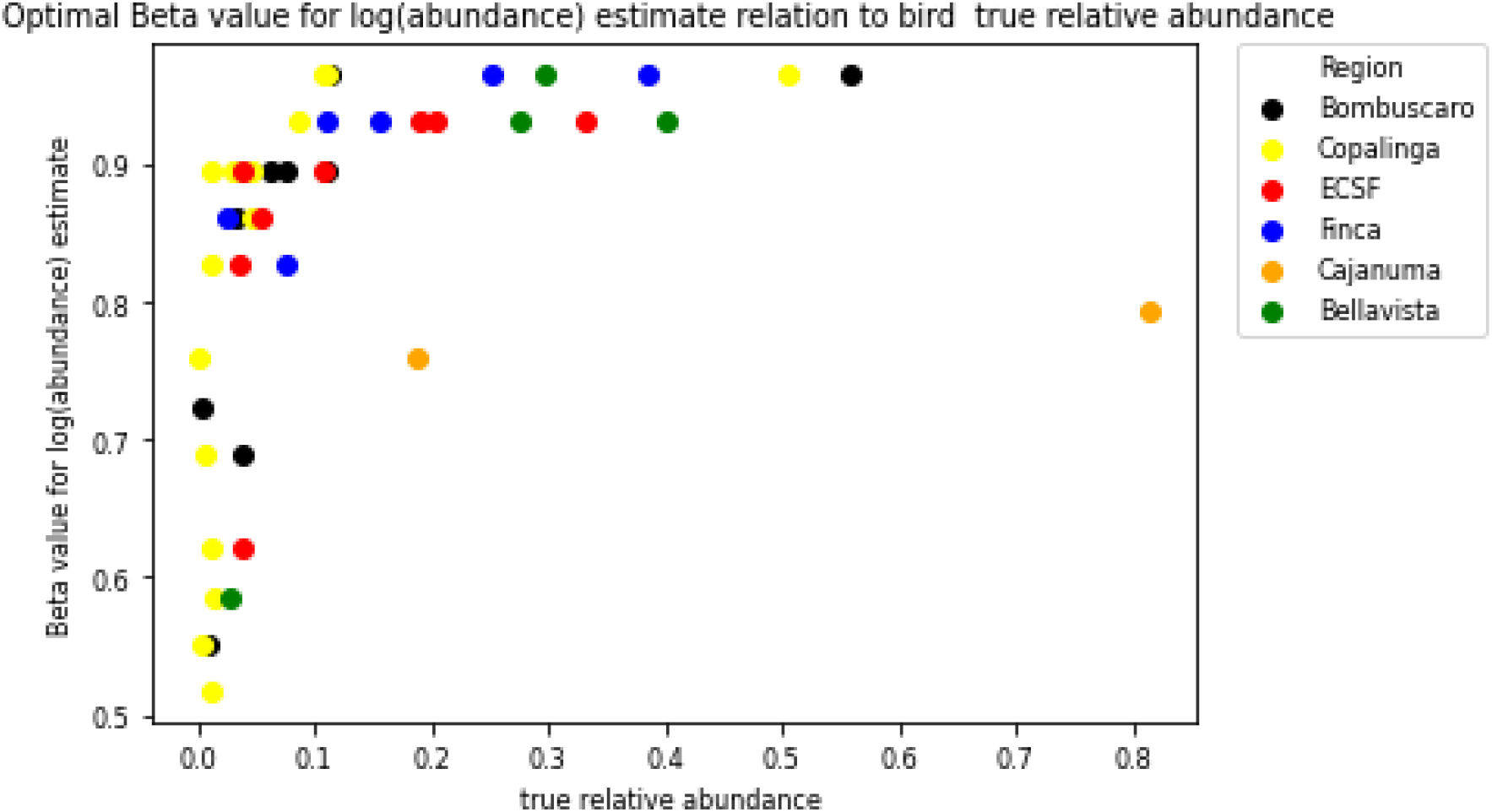
RMSE-minimizing value of *β* for the combined log abundance estimator 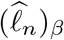, plotted against true relative abundance. Layout and interpretation as in Figure B6. Transect estimator uses 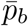 (case a).

**Figure B8:**
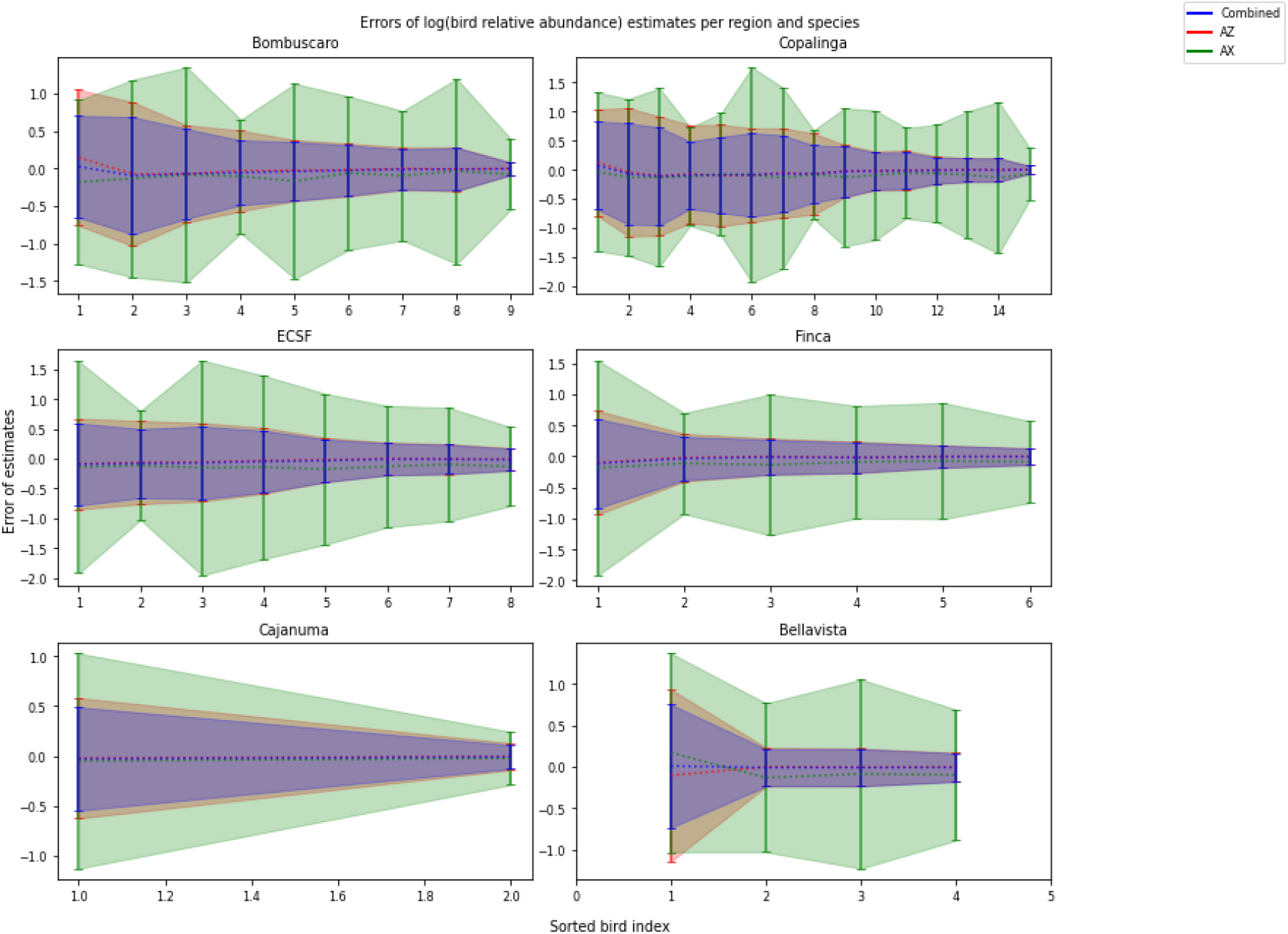
95% confidence intervals for the error of three log abundance estimators: 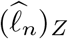 (labeled *AZ*), 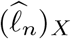 (labeled *AX*), and the combined estimator 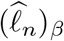. Results are shown for each bird family in each site. The *AX* and combined estimators use 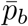 (case a). Families are ordered by increasing true relative abundance within each site panel.

## Supplement C: Estimator bias, variance, and RMSE using exact bird weight exponent *p*_*b,n*_ (case b)

This supplement contains figures corresponding to those in Supplement B, but computed using the exact bird weight exponent *p*_*b,n*_ for each family (case b as defined in Section 3.2.3). Comparison with Supplement B illustrates the improvement in estimator accuracy that results from precise knowledge of the weight-energy relationship.

**Figure C1:**
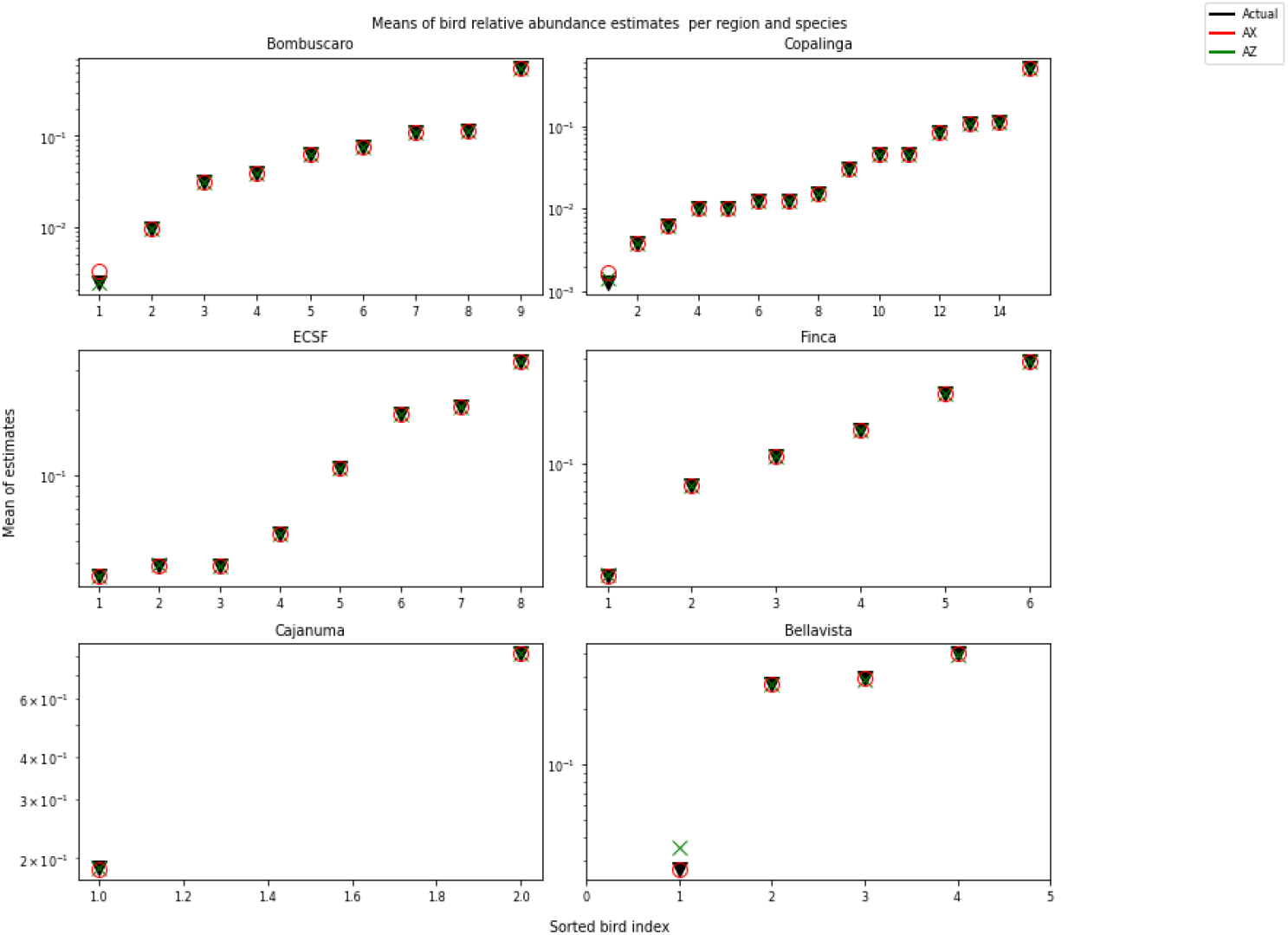
As Supplement B Figure B1, but with the transect-based estimator computed using the exact bird weight exponent *p*_*b,n*_ for each family (case b). Compared to Figure B1, the transect estimator shows reduced bias and tighter agreement with the true values, demonstrating the benefit of accurate knowledge of the weight-energy relationship.

**Figure C2:**
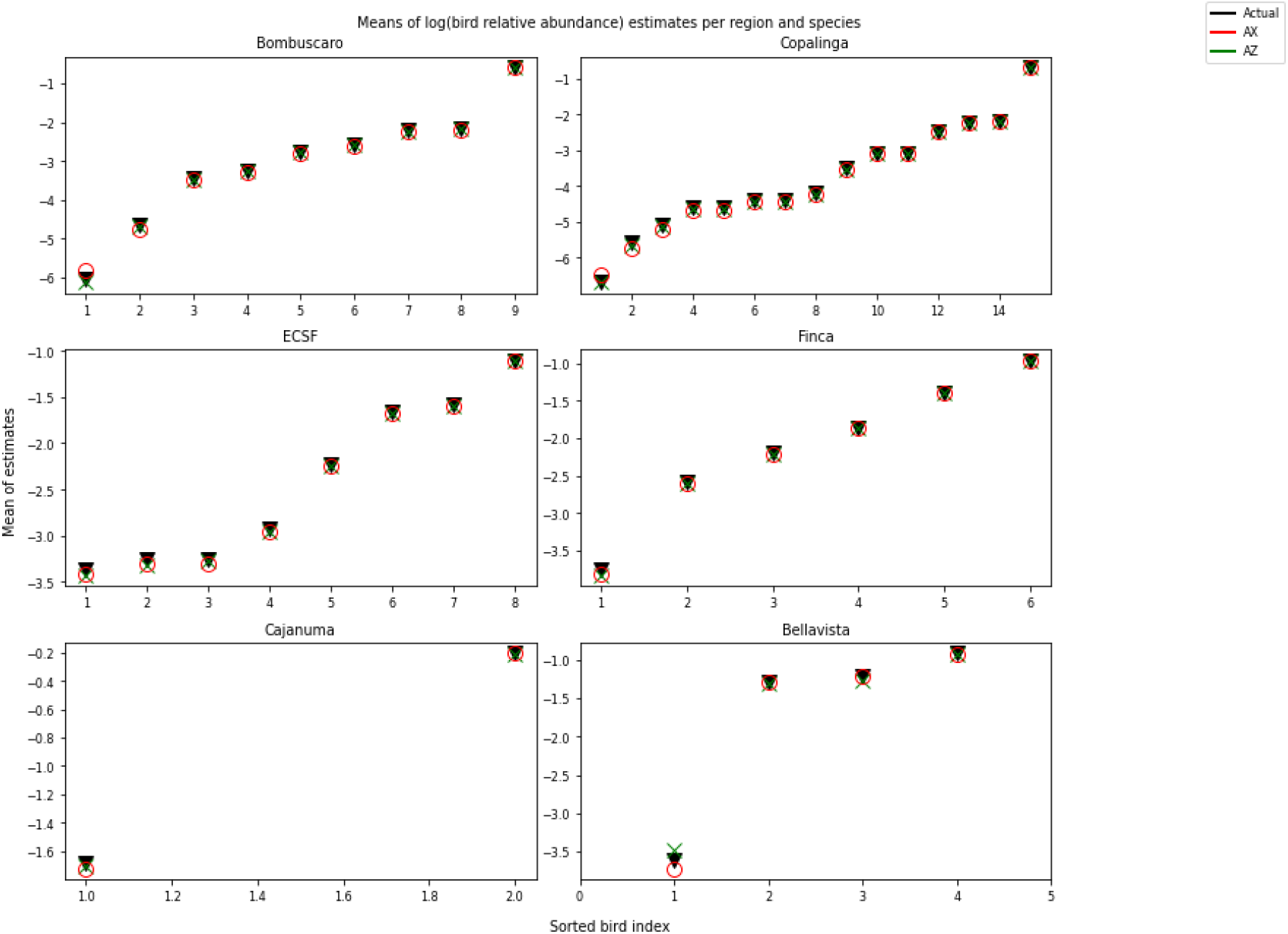
As Supplement B Figure B2, but with the transect-based log abundance estimator computed using the exact bird weight exponent *p*_*b,n*_ for each family (case b). Families are sorted as in Figure B1.

**Figure C3:**
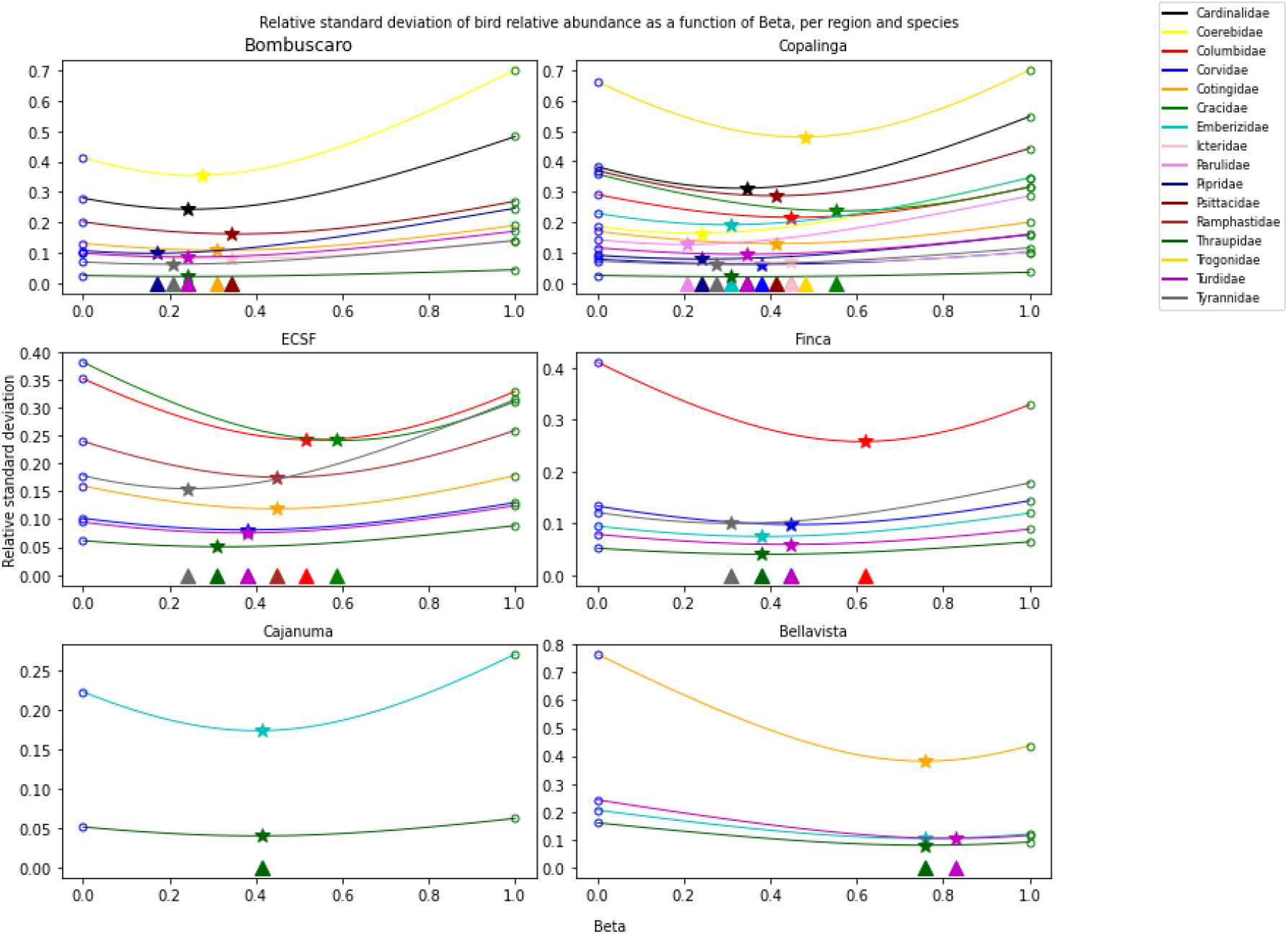
As main body Figure 4, but with the transect estimator using the exact bird weight exponent *p*_*b,n*_ (case b). The minimum standard deviation at *β* = 0 (pure transect estimator) is markedly lower than in Figure 4, and optimal *β* values shift toward lower values, reflecting the improved accuracy of the transect estimator when *p*_*b,n*_ is known precisely.

**Figure C4:**
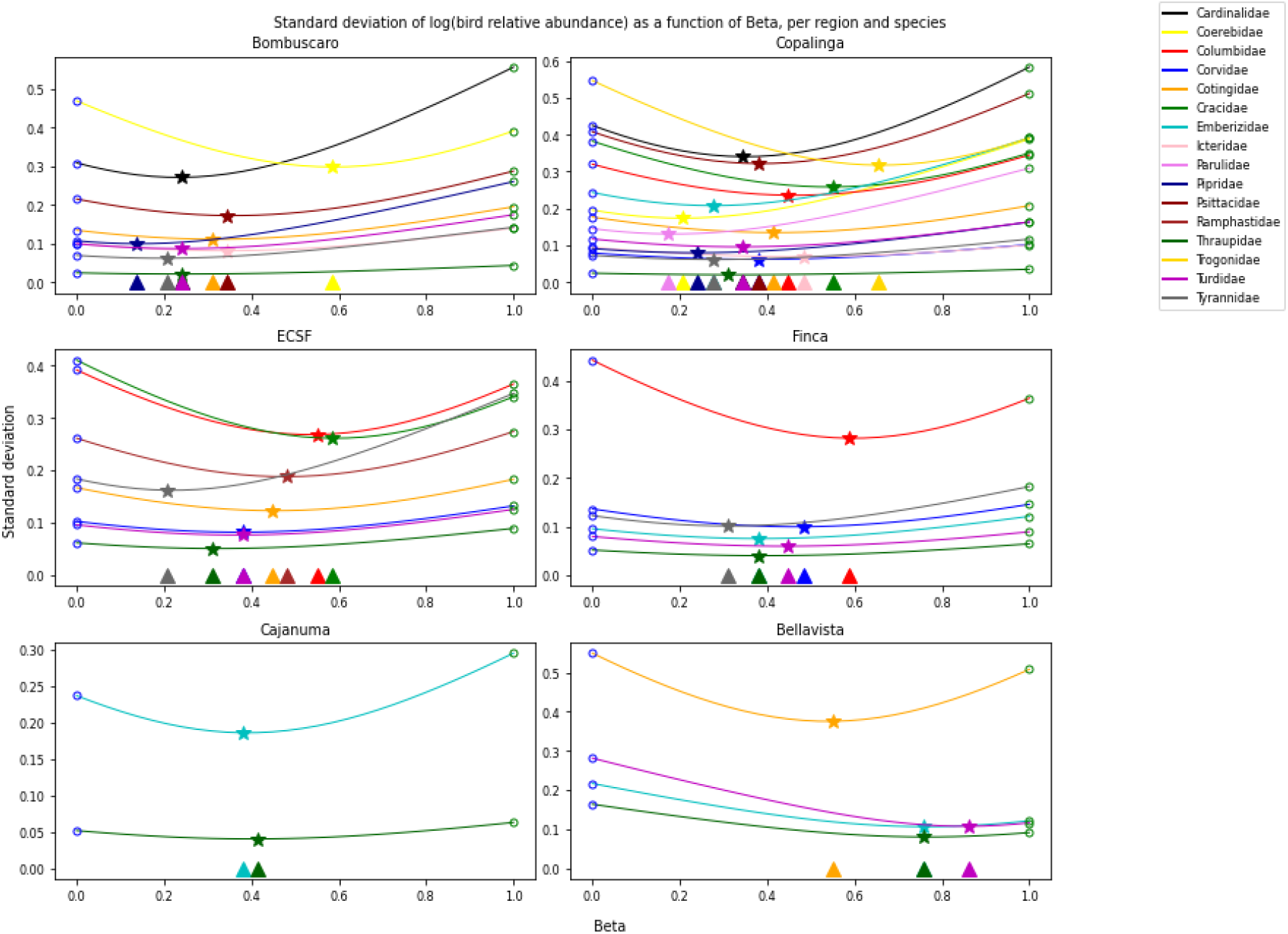
As Supplement B Figure B3, but with the transect log abundance estimator using the exact bird weight exponent *p*_*b,n*_ (case b). Optimal *β* values shift toward lower values compared to Figure B3, reflecting the improved transect estimator accuracy.

**Figure C5:**
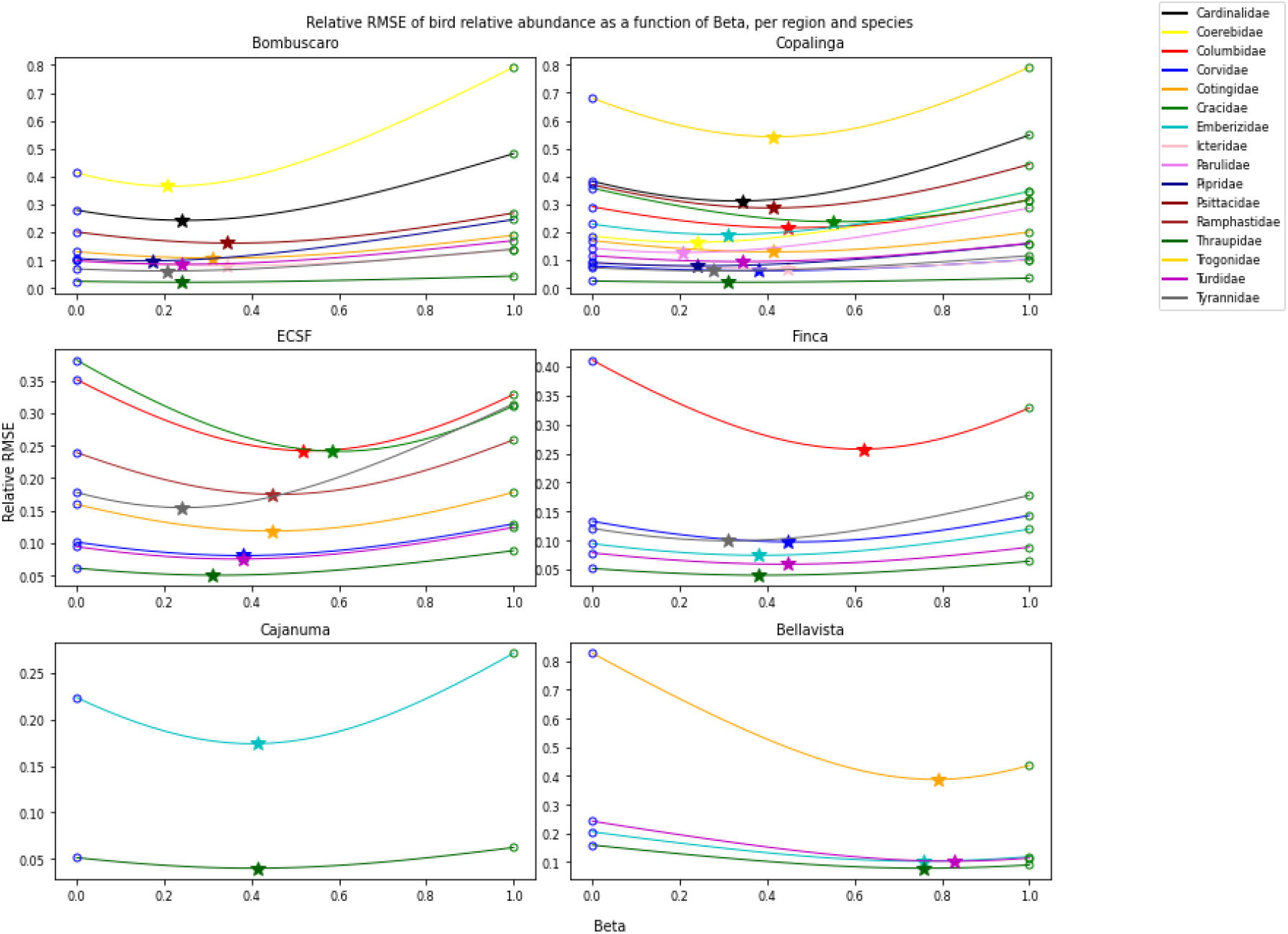
As Supplement B Figure B4, but with the transect estimator using the exact bird weight exponent *p*_*b,n*_ (case b). Minimum relative RMSE values are substantially lower than in Figure B4, confirming the advantage of precise knowledge of the weight-energy exponent.

**Figure C6:**
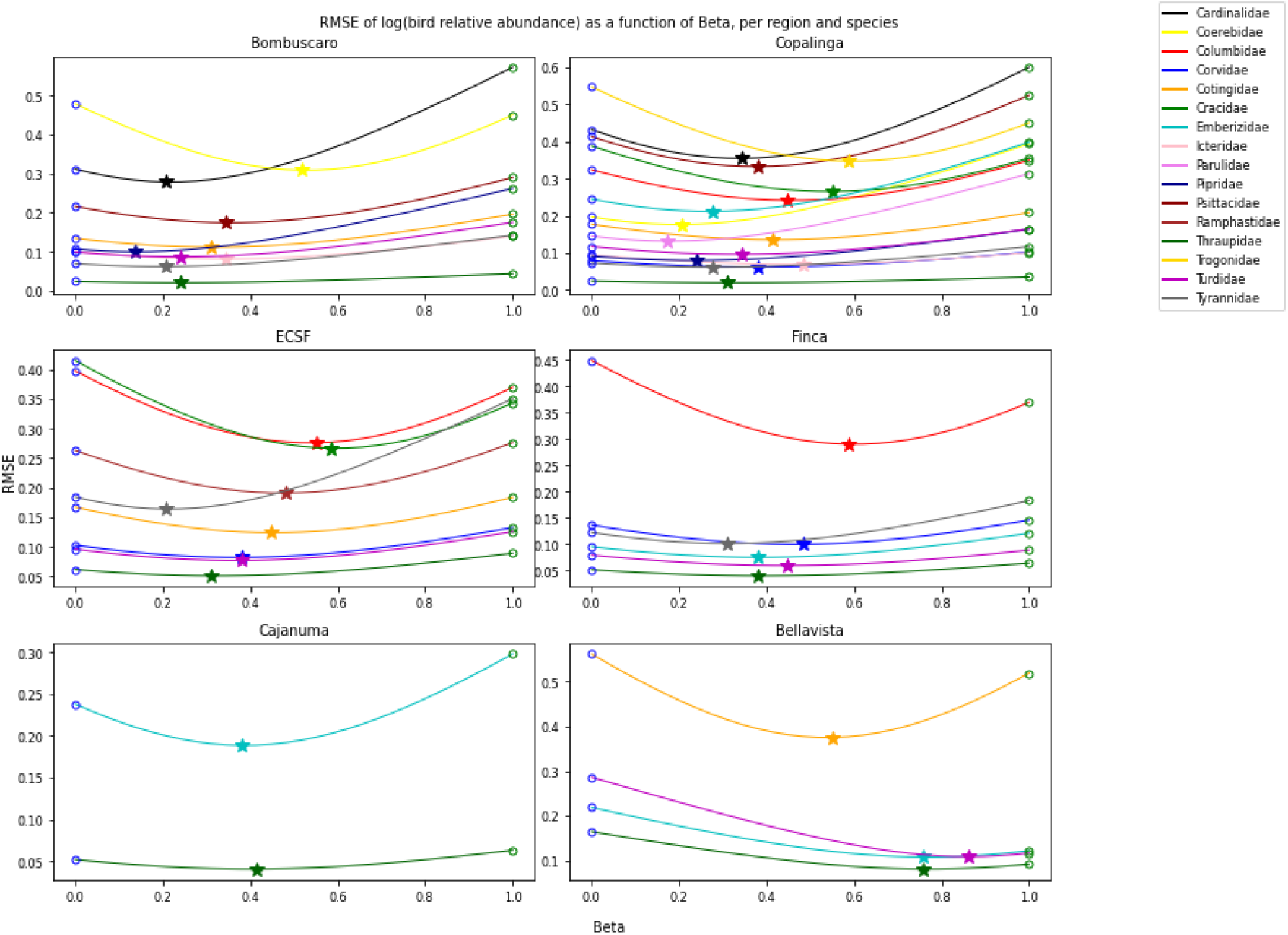
As Supplement B Figure B5, but with the transect log abundance estimator using the exact bird weight exponent *p*_*b,n*_ (case b). Minimum RMSE values are reduced relative to Figure B5.

**Figure C7:**
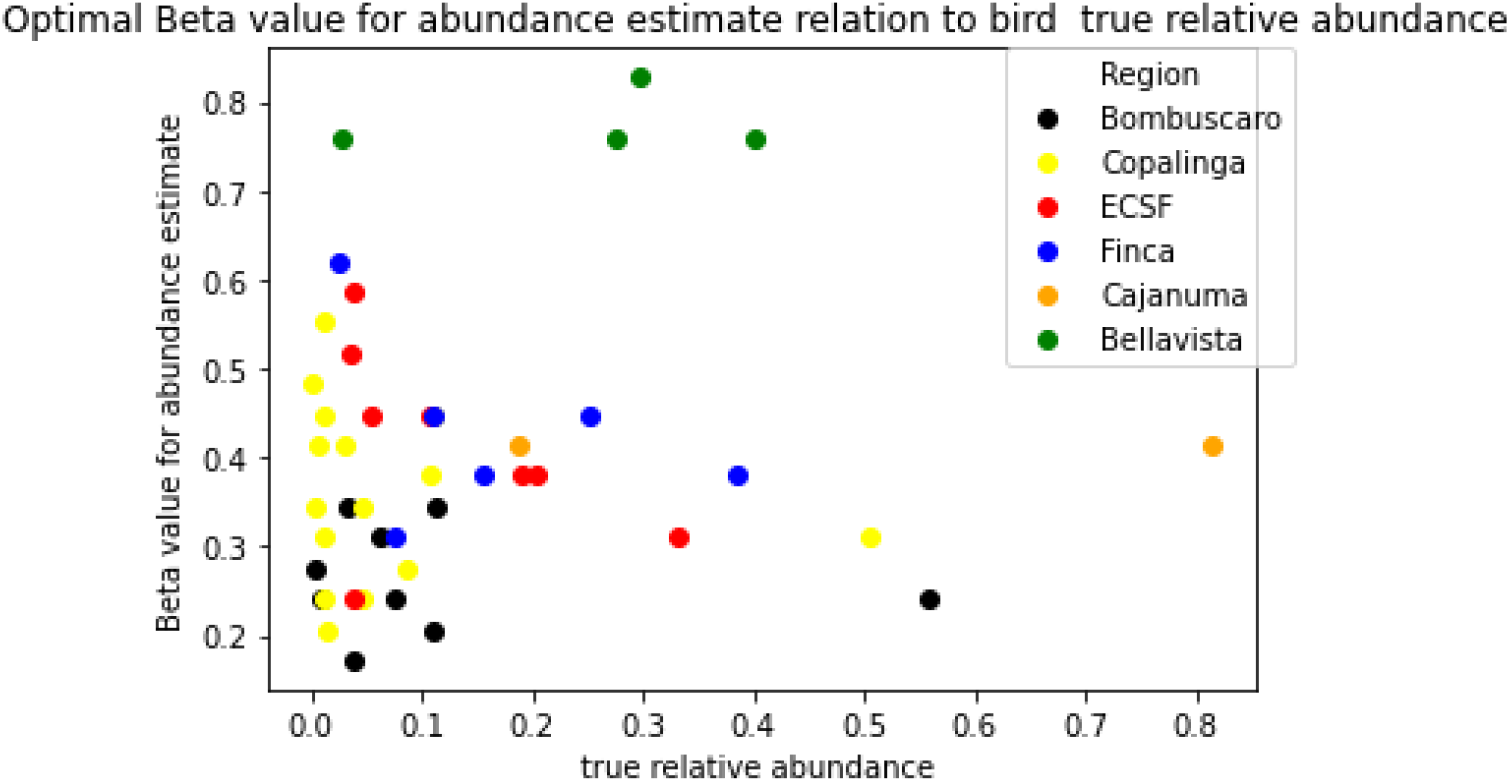
As Supplement B Figure B6, but with the transect estimator using the exact bird weight exponent *p*_*b,n*_ (case b). Optimal *β* values are generally lower than in Figure B6, indicating that more weight is appropriately assigned to the transect estimator when *p*_*b,n*_ is known precisely.

**Figure C8:**
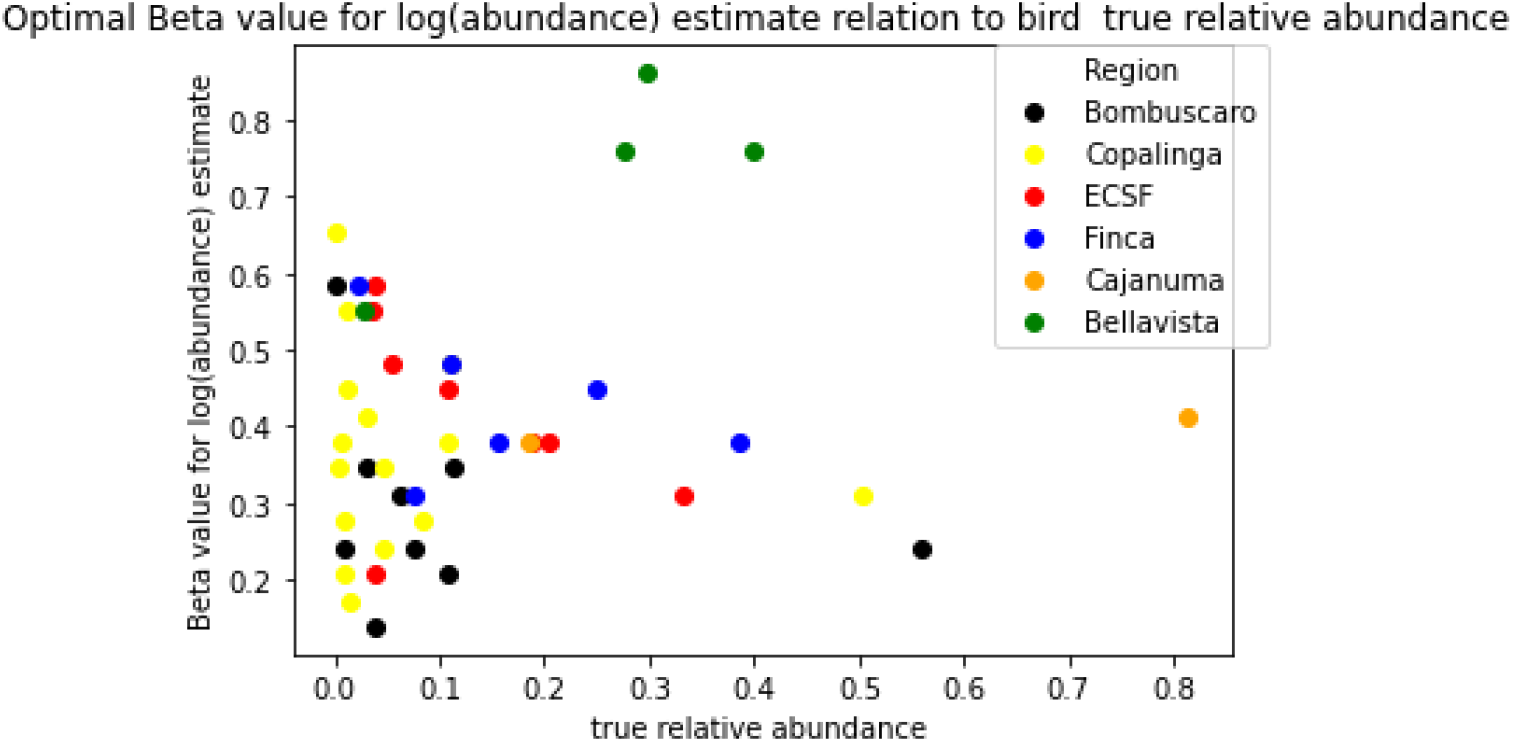
As Supplement B Figure B7, but with the transect log abundance estimator using the exact bird weight exponent *p*_*b,n*_ (case b). Layout and interpretation as in Figure C7.

**Figure C9:**
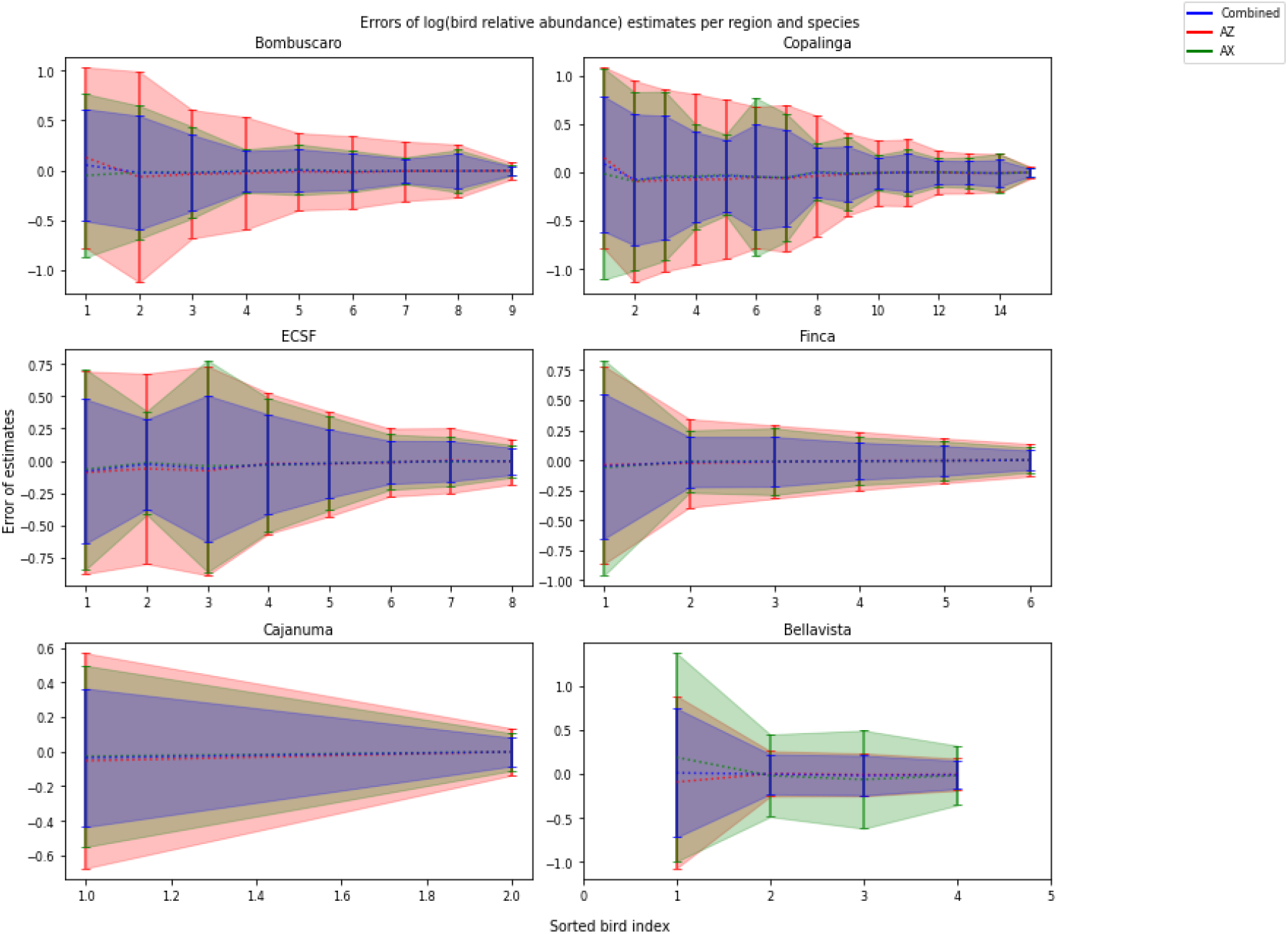
As Supplement B Figure B8, but with the transect-based and combined log abundance estimators using the exact bird weight exponent *p*_*b,n*_ (case b). Confidence intervals are substantially narrower for the transect and combined estimators compared to Figure B8.

## Notes

### Competing Interest Statement

The authors have declared no competing interest.

### Summary of Updates

Section 2 (Previous Studies) was expanded from a brief paragraph into three full paragraphs covering, respectively, the ecological importance of frugivore birds in tropical and subtropical systems, the advantages and limitations of point count and transect survey methods and the rationale for combining them, and the role of Bayesian and simulation-based statistical methods in improving abundance estimation reliability. Section 3.2.1 was expanded to include an ecological description of Podocarpus National Park's vegetation zones by elevation, an explicit characterization of fragmented sites as forest patches separated by cattle pasture and small-scale agriculture, and two new protocol paragraphs detailing the point count and transect survey procedures including observation window, detection radius, transect pace and width, and what was recorded per feeding event. A sentence was added to Section 3.2.2 defending the use of AIC over p-values for model comparison, and a complementary sentence was added to the Results clarifying that confidence intervals for p_f containing zero are equivalent to a significance test at the 5% level. A new Discussion section was added with four subsections addressing the biological interpretation of the fruit weight and bird weight findings, the practical implications of combining survey types for abundance estimation, the calibration of the bootstrapping confidence interval procedure, and the limitations of the current framework with suggestions for future work. All 31 figure captions were rewritten to be fully self-contained. The figures were reorganized so that 6 key figures appear in the main body and the remaining 25 are distributed across three supplements: Supplement A contains model fit diagnostic scatter plots for both the one- and two-coefficient models; Supplement B contains detailed estimator performance figures for case (a) using the mean bird weight exponent; and Supplement C contains corresponding figures for case (b) using the exact bird weight exponent. The Results text was revised throughout to direct readers to the appropriate supplement figures while preserving all original content and numerical results without abbreviation. Minor typographical errors in the original tables were also corrected.

